# Antagonistic Roles for MITF and TFE3 in Melanoma Plasticity

**DOI:** 10.1101/2024.07.11.603140

**Authors:** Jeremy Chang, Katelyn R. Campbell-Hanson, Marion Vanneste, Nicholas I. Bartschat, Ryan Nagel, Asdis K Arnadottir, Hong Nhung Vu, Collin Montgomery, Julius Yevdash, Jiarui Jiang, Ardith Bhinu, Annika Helverson, Michael D. Henry, Eiríkur Steingrímsson, Ronald J. Weigel, Robert A. Cornell, Colin Kenny

**Author notes:** Further information and requests for resources and reagents should be directed to and will be fulfilled by the lead contact, Colin Kenny.

## Abstract

Melanoma cells have the ability to switch from a melanocytic and proliferative state to a mesenchymal and invasive state and back again. This plasticity drives intra-tumoral heterogeneity, progression, and therapeutic resistance. Microphthalmia-associated Transcription Factor (MITF) promotes the melanocytic/proliferative phenotype, but factors that drive the mesenchymal/invasive phenotype and the mechanisms that effect the switch between cell states are unclear. Here, we identify the MITF paralog TFE3 and the non-canonical mTORC1 pathway as regulators of the mesenchymal state. We show that TFE3 expression drives the metastatic phenotype in melanoma cell lines and tumors. Deletion of TFE3 in MITF-low melanoma cell lines suppresses their ability to migrate and metastasize. Further, MITF suppresses the mesenchymal phenotype by directly or indirectly activating expression of *FNIP1, FNIP2,* and *FLCN*, which encode components of the non-canonical mTORC1 pathway, thereby promoting cytoplasmic retention and lysosome-mediated degradation of TFE3. These findings highlight a molecular pathway controlling melanoma plasticity and invasiveness.

## Introduction

Cellular plasticity is a fundamental aspect of tumor biology that drives cancer progression and therapeutic resistance. Plasticity promotes aggressive tumor cell states through the activation of transcriptional programs associated with stemness and dedifferentiation (reviewed in ^1,2^). A knowledge of cancer plasticity and intra-tumor heterogeneity is central to understanding the cellular mechanisms that govern the invasiveness of tumors and their resistance to therapy (reviewed in ^3^). The cancer stem cell (CSC) model, according to which a few cancer cells have self-renewing capacities, provides an explanation for how tumor heterogeneity and hierarchy can be maintained. CSCs exhibit high plasticity, switching between tumor-initiating stem-cell like, rapidly-proliferating, and differentiated states through environmental, epigenetic, and transcriptional stimuli (reviewed in ^1,2,3^). Targeting cellular plasticity is a promising therapeutic approach but will require a deeper understanding of the mechanisms that are critical for these cell state transitions. Outstanding questions related to these mechanisms include: do core programs exist that define plasticity across cancer types? What is the relationship between plasticity and proliferative potential? What are the consequences of ablating high-plasticity cell states for tumor progression or response to therapy?

Cutaneous melanoma, the most lethal cutaneous cancer, is characterized by cellular plasticity. Even within a single melanoma tumor, individual cells can be distinct phenotypically, transcriptionally, and epigenetically, as well as with respect to their sensitivity to therapy ^4–8^. Analyses of genome-wide expression profiles in melanoma cell lines and tumors have detected a common trend based on activity of the microphthalmia-associated transcription factor (MITF), a master regulator of melanocyte differentiation ^9,10^. *Mitf*-null mice lack melanocytes, are deaf, and have unpigmented retinal pigment epithelia (reviewed in ^11^). In melanoma, high MITF activity correlates with a relatively proliferative and pigmented phenotype, and with high expression of genes promoting these qualities, while low MITF activity correlates with increased invasive behavior and expression of genes promoting epithelial-mesenchymal transition (EMT), invasion, and stem-cell-like or neural-crest-cell-like properties ^4,6,8,10^. Low MITF activity is also correlated with drug resistance ^12^. Melanoma tumor cells can transition between these phenotypes (phenotype switching) ^6,7,10^. Switching from the MITF-high to MITF-low state is characterized by re-emergence of a transcriptional state resembling that of the neural crest, the embryonic precursor population from which melanocytes are derived ^13^, and is critical for the initiation of melanoma ^4,6,7,10,13–16^. MITF activity and expression are in part regulated by nutrient availability ^17,18^, hypoxia ^19,20^, and inflammatory signaling ^18^, suggesting such mechanisms will influence MITF-dependent cell states. Elucidating the transcriptional mechanisms that govern switching across these phenotypes may facilitate the design of therapies to target invasive and therapy-resistant melanoma.

Understanding mechanisms of phenotype switching necessarily requires an understanding of how the two major phenotypes are controlled. Genomic occupancy experiments, such as ChIP-Seq or CUT&RUN-seq, in MITF-high melanoma, and RNA sequencing of such cells after *MITF* has been deleted, indicated that MITF directly activates a cohort of genes promoting pigmentation and proliferation ^11,21–23^. These findings explain why in MITF-low melanoma expression of such genes is low, but not why expression of genes promoting EMT, invasion, dedifferentiation and stemness genes is high. Interestingly, many of the latter types of genes are flanked by MITF peaks in an MITF-high melanoma cell line, which raises the possibility that MITF directly represses the expression of such genes ^22^. To test this possibility, we initiated the present study. We profiled MITF binding and chromatin marks reflective of active enhancers in SKMEL28 melanoma cells that were wild type or that harbored loss-of-function mutations in the *MITF* gene (i.e., *MITF*-*WT* vs. *MITF*-*KO*). The results argued against MITF functioning as a transcriptional repressor, at least when binding as a homodimer. Instead, they supported a TFE3-dependent mechanism by which characteristic genes are upregulated in MITF-low melanoma, and an FNIP2-dependent mechanism by which TFE3 is suppressed in MITF-high melanoma. We tested these mechanisms in a panel of MITF-high and MITF-low patient-derived xenograft (PDX) and commercial melanoma cell lines.

## Results

### Chromatin profiling suggests that MITF directly activates but does not directly repress chromatin and gene expression

We recently conducted CUT&RUN in SKMEL28 cells using a commercial antibody generated against a peptide within MITF ^22,23^. However, slight variants on this peptide are present within the MITF paralogs TFEB, TFE3, and TFEC (**Figure S1A**), raising the possibility that a subset of the peaks reflect binding by an MITF paralog instead of by MITF. We reasoned that peaks present in wild-type (WT) cells but absent from *MITF* knockout (KO) cells would most likely represent binding by MITF homodimers (although they could also be binding by a heterodimer of MITF and a paralog) and that peaks present in both wild-type (WT) cells and *MITF* knockout (KO) cells would represent binding of a paralog homodimer or a heterodimer. We reanalyzed our previously-published anti-MITF CUT&RUN data from SKMEL28 cells ^22,23^ and conducted anti-MITF CUT&RUN data in two previously-generated SKMEL28 clones lacking functional *MITF* alleles: Δ*MITF-X6* and Δ*MITF-X2* (hereafter, *MITF*-*KO*, D6 or D2 cells) ^22^. *MITF*-*KO* D6 cells lack MITF protein detectable on Western blot, while *MITF*-*KO*, D2 cells retain a low molecular weight protein recognized by anti-MITF antibody^22^. Approximately 84% of anti-MITF peaks identified in D6 cells overlapped with those in D2 cells (**Figure S2A**).

Among the 46,519 anti-MITF peaks in WT SKMEL28 cells, only 10% (4,593) were absent in the *MITF-KO* D6 cells; hereafter we will refer to this subset as true MITF peaks (**Figure 1A**, blue rectangle). Approximately 75% of the peaks in *MITF*-KO cells (34,623) were shared with *MITF*-WT cells, suggesting that an MITF paralog, such as TFE3, TFEB or TFEC, occupies these sites in both genotypes or specifically occupies them in the absence of MITF (**Figure 1A**, grey rectangle). The latter possibility suggests potential competition between paralogs for binding at these loci. Alternatively, these sites may reflect heterodimers of MITF and its paralogs in WT cells that become homodimers or heterodimers of TFE3, TFEB or TFEC in *MITF*-KO cells. We refer to this subset as persistent-paralog peaks. Interestingly, there were 6,943 anti-MITF peaks unique to *MITF-KO* cells, apparently reflecting binding of a paralog gained upon the loss of MITF; we designated this subset as gained-paralog peaks (**Figure 1A**, yellow rectangle).

**Figure 1:**
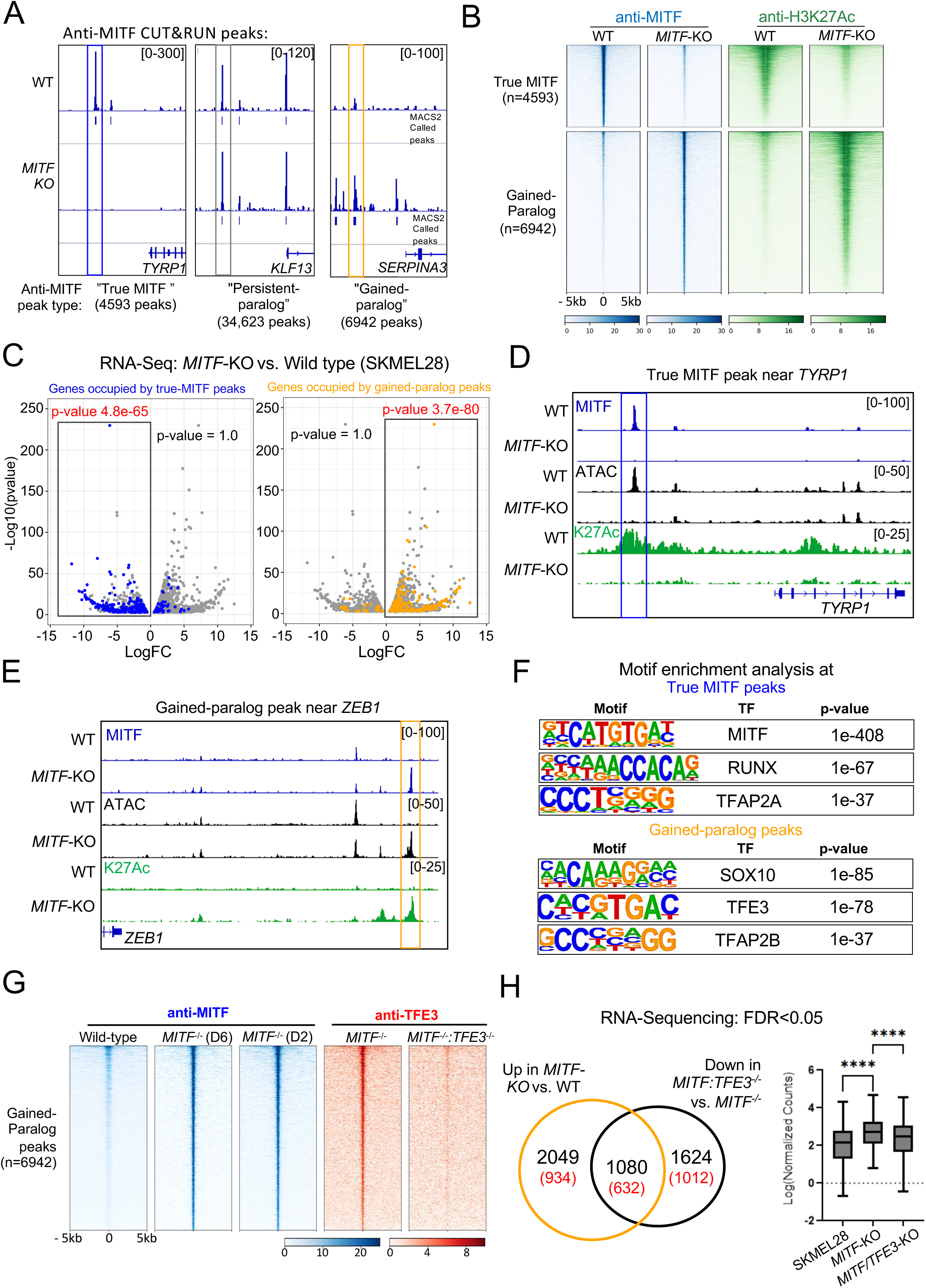
MITF inhibits TFE3 occupancy at thousands of genomic loci in SKMEL28 cell lines. (**A**) Screenshot of IGV genome browser (GRCH37/hg19) visualization of bigwig files generated from anti-MITF CUT&RUN sequencing in *MITF*-wild-type (WT) or *MITF-KO* (D6) cells. Three groups of anti-MITF peaks (true MITF, persistent-paralog, and gained paralog) were identified based on the presence or absence of an anti-MITF peak in *MITF-KO* cells. The number of each peak type is indicated. Peaks were called using MACS2 software with FDR <0.05, with IgG serving as background control. (**B**) Density heatmap of true MITF peaks (upper panels) and gained-paralog peaks (lower panels), showing anti-MITF (blue) and anti-H3K27Ac (green) CUT&RUN signal in *MITF-WT* and *MITF-KO* cells, with peaks sorted on anti-MITF-signal in *MITF-WT* cells. (**C**) Volcano plot showing 2,144 differentially expressed genes (DEGs) with an adjusted p-value < 0.05 in D6 versus EV-SKMEL28 cells. RNA-Seq data was re-analyzed from Dilshat et al (2021) ^22^, and DEGs were overlapped with genes flanked by true MITF peaks and gained paralog peaks. Peaks were assigned to genes based on proximity (±100 kb from the gene’s transcription start site [TSS]). A hypergeometric analysis was performed to determine the association of true MITF peaks and gained paralog peaks with DEGs. p-values are labeled, and significant associations are indicated in red text. (**D-E**) Screenshots of IGV browser illustrating (**D**) a true MITF peak near *TYRP1*, and (**E**) a gained-paralog peak near *ZEB1*. Additional tracks include ATAC-Seq (black), and anti- H3K27Ac CUT&RUN (green) in *MITF-WT* and *MITF-KO* cells. (**F**) Enrichment of transcription-factor motifs at true MITF peaks (upper panel) and at gained-paralog peaks (lower panel), as assessed using HOMER ^24^. (**G**) Density heatmap of anti-MITF CUT&RUN-seq in *MITF-WT* and *MITF-KO* cells (both D6 and D2 lines), and of anti-TFE3 CUT&RUN-seq in *MITF-KO* and double *MITF/TFE3- KO* cells (*D2/TFE3-KO*), at gained-paralog peaks. (**H**) Venn diagram illustrating the overlap of genes that are upregulated in *MITF-KO* (D2) vs *MITF-WT* cells (depicted by the orange circle), and the genes that are downregulated in double *TFE3/MITF-KO* (*D2/TFE3*-KO) vs. *MITF-KO* (D2) cells (depicted by the black circle). Gene numbers in red parenthesis illustrate TFE3 occupied genes (peak within 100kb of a genes TSS) in *MITF*-KO cells. Box-plot visualization of Log normalized counts of the overlapping genes (n=632). Student’s t-test; ****p-value <0.0001.

Across duplicate CUT&RUN experiments, true MITF peaks were highly correlated (r = 0.97, Pearson correlation), and persistent-paralog peaks and gained-paralog peaks slightly less so (r = 0.84 and 0.88, respectively) (**Figure S2B-C**).

To determine the functions of these subsets of anti-MITF peaks, we assessed chromatin accessibility and activity in *MITF-WT* and *MITF-KO* cells. We performed ATAC-seq and CUT&RUN using antibodies targeting post-translational marks on histone H3, including the activation marks chromatin lysine 27 acetylation (H3K27Ac) and lysine 4 tri- methylation (H3K4Me3) and the repressive mark lysine 27 tri-methylation (H3K27Me3). Active chromatin was positive for ATAC-seq and H3K27Ac signals and lacking H3K27Me3 signal. All true MITF peaks were located within active chromatin in *MITF-WT* cells (**Figure S3A**), and at 92% of them the H3K27Ac signal was reduced in *MITF-KO* D6 cells (**Figure 1B and Figure S3A-B**). To test the notion that this subset corresponds to enhancers directly activated by MITF we re-analyzed RNA seq on SKMEL28 *MITF-WT* and *MITF-KO* D6 cells ^22^. We found that genes associated with this subset of true MITF peaks almost all had higher expression in *MITF-WT* than in *MITF-KO* cells (hypergeometric analysis p-value 4.8e-65; **Figure 1C**: blue dots correlate with downregulated genes, **and Table S1**), and, as expected, these genes were enriched for gene ontology (GO) terms related to melanocyte development (e.g., *DCT*, *OCA2*, *GPR143*, *TRPM1* and *TYRP1*) (**Figure S3C**). An example of an MITF-activated enhancer near *TYRP1*, a gene involved in melanin synthesis, is shown in **Figure 1D**. At the remaining 8% of true MITF peaks, the levels of H3K27Ac signals were equivalent in the *MITF-WT* and *MITF-KO* cells (**Figure S3A-B**). Importantly, at no true MITF peaks was there the expected signature of enhancers directly repressed by MITF, i.e., higher levels of ATAC-seq and H3K27Ac signals in *MITF-KO* relative to in *MITF-WT* cells (**Figure 1B** and **Figure S3A-B**). In summary these results indicate that, when binding as a homodimer, MITF activates chromatin but does not inactivate it.

At persistent paralog peaks, the anti-MITF signal was equivalent in *MITF-WT* and *MITF-KO* cells (**Figure S3A**) and the chromatin was active in both *MITF-WT* cells and *MITF-KO* cells. Although these peaks were not the focus of this study, we note that at 27% of persistent-paralog peaks, the ATAC and H3K27Ac signals were higher in *MITF-KO* cells than in *MITF-WT* cells, and that this subset was associated with genes whose expression was higher in *MITF-KO* cells than in *MITF-WT* cells (hypergeometric analysis p-value 1.02e-73). These observations suggest that certain enhancers are activated by a heterodimer of MITF and an MITF paralog, but are activated more effectively by paralog(s) binding without MITF.

Finally, gained-paralog peaks were located within chromatin that was inactive in *MITF-WT* cells but active in *MITF-KO* cells (**Figure 1B, 1E, Figure S3A-B**). The genes associated with gained-paralog peaks often had higher expression in *MITF-KO* cells than in *MITF-WT* cells (hypergeometric analysis p-value 3.7e-80; **Figure 1C**, yellow dots correlate with genes upregulated in *MITF-KO* cells, **Table S1**). Interestingly, genes associated with gained-paralog peaks and with higher expression in *MITF-KO* cells than in *MITF-WT* cells were enriched for GO terms including cell migration, cell motility, extracellular remodeling, and epithelial to mesenchymal transition (EMT) (e.g., *TGFA, ITGA2, MMP15, FGF1, SERPINA3, ZEB1*) (**Figure S3D**). An example of a gained- paralog peak near *ZEB1*, whose expression is associated with EMT, is shown in **Figure 1E**. These findings indicate that gained-paralog peaks activate enhancers near genes associated with migratory behavior.

### TFE3 binds at gained-paralog peaks

To determine the identity of the MITF paralog that binds at gained-paralog peaks we performed a motif enrichment analysis of chromatin elements underlying these peaks using the Hypergeometric Optimization of Motif EnRichment (HOMER) software ^24^. As expected, elements underlying true MITF peaks were strongly enriched for the binding sites of MITF and of its co-factor TFAP2A (**Figure 1F**) ^23^. Interestingly, chromatin elements underlying gained-paralog peaks that were associated with genes with higher expression in *MITF-KO* cells than in *MITF-WT* cells were most strongly enriched for the binding site of the MITF paralog TFE3. They were also enriched for the binding site of SOX10 and TFAP2B, suggesting potential co-factors for TFE3 (**Figure 1F**). Next, we used the anti-MITF antibody to immuno-precipitate protein from lysates of the *MITF-WT* and *MITF-KO* cells. On Western blots of protein immunoprecipitated from cells of both genotypes, anti-TFE3 antibody recognized bands at the appropriate size (∼72–82 kDa) (**Figure S4A-C**). We cut out these bands, extracted protein, subjected it to mass spectrometry, and detected peptides derived unambiguously from TFE3 (**Figure S4D**). Next, we deleted the *TFE3* gene from *MITF-KO* D2 cells then conducted anti-TFE3 and anti-MITF CUT&RUN in *MITF-KO* D2 cells and *MITF/TFE3* double-KO cells. (We also deleted TFE3 in *MITF-KO* D6 cells but found that the resulting *MITF/TFE3* double-KO survived for fewer passages than those created from *MITF-KO* D2 cells). In *MITF-KO* cells an anti-TFE3 peak overlapped all of the gained-paralog subset of anti-MITF peaks (**Figure 1G**), while in *MITF/TFE3* double-KO cells, all of these anti-TFE3 peaks were absent (**Figure 1G**). Moreover, the anti-MITF CUT&RUN signal at gained-paralog peaks was strongly reduced or lost in *MITF/TFE3* double-KO cells, further supporting that the anti-MITF antibody cross-reacts with TFE3 (examples shown in **Figure S4E**). The anti-MITF CUT&RUN signal at persistent paralog peaks was ∼50% reduced in *MITF/TFE3* double-KO cells, indicating that a different MITF paralog, i.e., TFEB or TFEC, binds at them, perhaps only in the absence of TFE3 (**Figure S4F**). We refer to gained-paralog peaks as gained-TFE3 peaks.

Finally, we amplified a DNA element underlying a gained-TFE3 peak 10 kb upstream of the transcription start site of *SERPINA3* (**Figure S5A**), a gene whose expression is higher in *MITF*-KO (D2 and D6) cells than in *MITF*-WT cells, making this element a candidate for an enhancer activated by TFE3. We engineered it into a reporter construct with a minimal promoter and the firefly luciferase gene, and transfected the construct into *MITF-WT*, *MITF*-*KO* D2, and *MITF* D2*/TFE3* double-*KO* cells. As expected, reporter activity from this construct was highest in the *MITF*-*KO* cell lines (**Figure S5A’**).

Moreover, when we deleted the predicted TFE3 binding sites from the element, its enhancer activity was no longer higher in *MITF*-*KO* D2 cells than in WT or *MITF* D2*/TFE3* double-*KO* cells (**Figure S5A’**). Together these results indicate that in *MITF-KO* cells, TFE3 binds and activates enhancers that are silent in *MITF-WT* cells and drives a unique transcriptional program.

### TFE3 activates expression of genes in the MITF-low transcriptional profile

We predicted that a subset of genes with higher expression in *MITF*-*KO* cells compared to in *MITF*-*WT* cells would be directly activated by TFE3. To test this, we performed RNA sequencing on *MITF*-*WT*, *MITF*-*KO* D2, and *MITF* D2*/TFE3* double-*KO* cells. Indeed, approximately 35% (1080/3129) of genes with elevated expression in *MITF*-*KO* cells relative to in *MITF*-*WT* cells had reduced expression in *MITF/TFE3* double-*KO* cells relative to in *MITF*-*KO* cells (**Figure 1H, Table S2-S3**), and the majority (58%: 632/1080) of these genes were flanked by anti-TFE3 peaks within 100 kb of their transcription start site, suggesting their expression is directly activated by TFE3. This apparently directly- TFE3-activated set of genes was enriched for GO terms related to cell migration (**Figure S5B**), cell differentiation, and angiogenesis (**Figure S5C**). In contrast, the remaining set of 2049 genes with higher expression in *MITF-KO* cells compared to in *MITF-WT* cells but with unchanged expression in *MITF/TFE3* double-KO relative to in *MITF-KO* cells was not enriched for EMT-related GO terms (**Figure S5D**), emphasizing the importance of TFE3 in promoting EMT and invasion in this cell type lacking MITF. Interestingly, within this latter set, 45% (934/2049) of genes was nonetheless flanked by anti-TFE3 peaks. This subset included *SOX2, TBX3, TBX6, PRDM12, FOXD2*, *FOXD1, FOXD3*, several members of the KLF transcription factor and WNT signaling pathways (**Table S4**). Notably, many of these genes were found to be regulated by TFE3 in naïve human embryonic stem cells (hESCs) ^25^. These results raise the possibility that TFE3 facilitates activation of genes promoting a stem-cell like state in melanoma as in hESCs, but is not necessary to maintain their expression following loss of TFE3. In summary, in an MITF-high cell line depleted of MITF, TFE3 drives the expression of genes associated with cell migration and stem-cell-like features.

Melanoma cell lines with low MITF activity (i.e., MITF-low) are characterized by expression of genes promoting cell invasion and stem-cell features ^4–8^. Given the results discussed above, we predicted that in MITF-low melanoma many such genes would be activated by TFE3. The invasive A375 human melanoma cell line expresses far lower levels of *MITF* than the proliferative *MITF-WT* SKMEL28 human melanoma cell line (**Figure S5E**). To identify active enhancers in A375 cells we assessed published ATAC-seq data and our H3K27Ac and H3K9Ac CUT&RUN-seq data from A375 cells and discovered that most (65%) gained-TFE3 peaks in *MITF-KO* cells overlapped the position of active enhancers in A375 cells (**Figure 2A**). Conversely, virtually all (98%) enhancers that are bound and activated by true MITF peaks in SKMEL28 cells are inactive in A375 cells (**Figure 2A**). We next examined ATAC-Seq data from three additional melanoma cell lines previously classified as melanocytic, and two each classified as intermediate or mesenchymal, based on their expression of genes characteristic of pigmentation and mesenchyme ^4^. In the melanocytic cell lines, the ATAC-seq signal was high at enhancers activated by true-MITF peaks in *MITF-WT* SKMEL28 cells (**Figure 2B**, MM001), and in mesenchymal cells it was high at enhancers activated by gained-TFE3 peaks in the *MITF-KO* (D2 and D6) cells (**Figure 2B**, MM029 and MM099). In the intermediate cell lines, the ATAC-seq signal was moderately high at both enhancer types (**Figure 2B**, MM074). Examples of ATAC-seq peaks in each category of melanoma cell lines near the MITF-activated gene *GPR143* and the TFE3- activated gene *ZEB1* are shown in **Figure 2C**. In summary, many enhancers that are activated upon deletion of *MITF* from an MITF-high cell line are also active in MITF-low melanoma cell lines.

**Figure 2:**
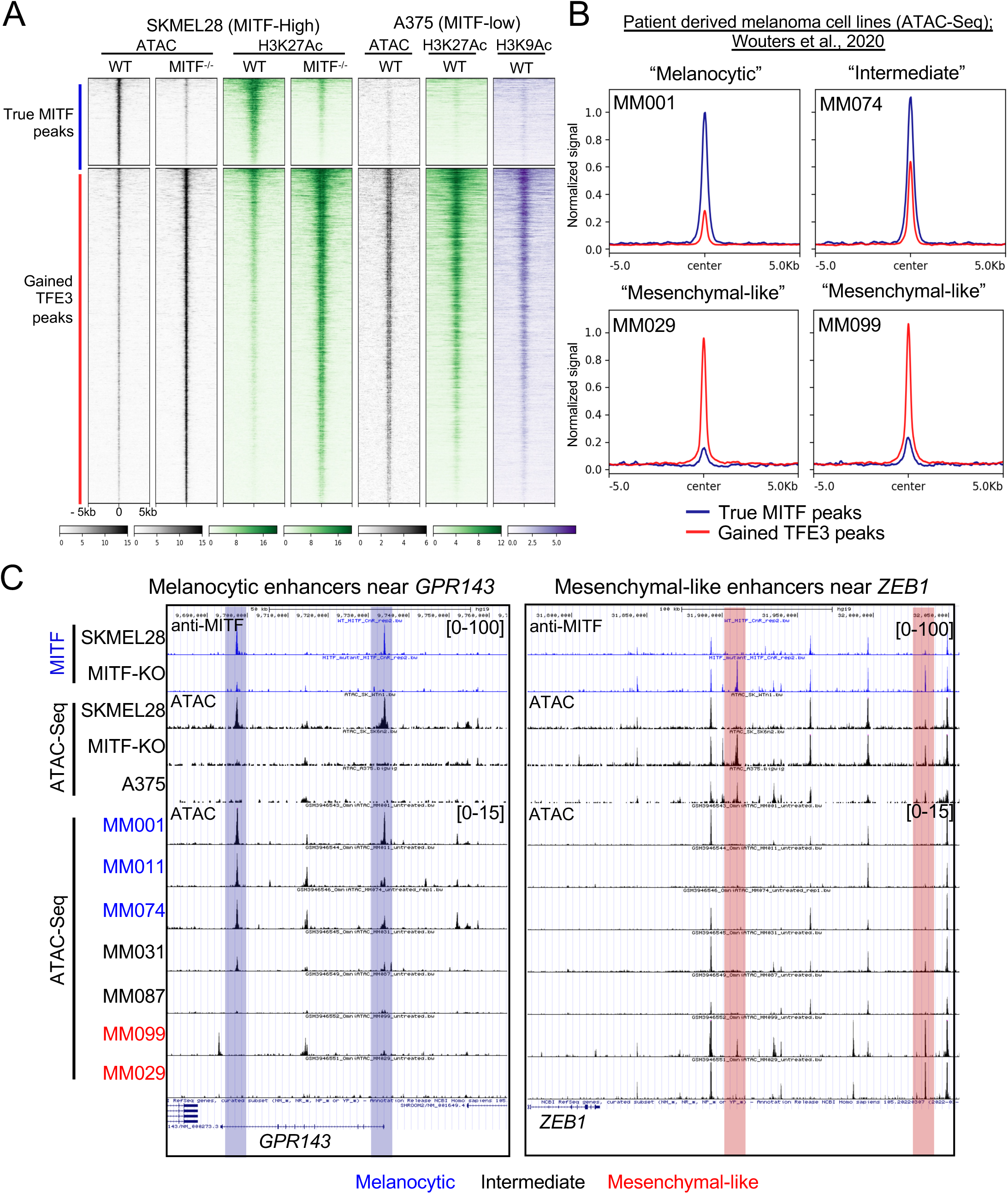
Gained TFE3 peaks in *MITF-KO* cells overlap with active enhancers in A375 and patient-derived "mesenchymal-like" cell lines, distinguishing them from true MITF bound "melanocytic" enhancers. (**A**) Density heatmap depicting ATAC-Seq (black) and anti-H3K27Ac (green) CUT&RUN-seq in *MITF-WT* and *MITF-KO* SKMEL28 cells, alongside ATAC-Seq (black), anti-H3K27Ac (green), and H3K9Ac (purple) CUT&RUN-seq in A375 cells. The density heatmap is ordered by true MITF peaks (upper panel) and gained-TFE3 peaks (lower panel), as identified in *MITF-WT* and *MITF-KO* SKMEL28 cells, respectively. (**B**) Summary plots illustrating normalized ATAC-Seq read counts at true MITF (blue line) and gained-TFE3 peaks (red line) for “melanocytic” (MM001), “intermediate” (MM074), and two “mesenchymal-like” (MM029, MM099) human patient-derived melanoma cell lines ^4^. (**C**) Screenshot of IGV genome browser (GRCH37/hg19) visualization of enhancers activated by true MITFs (blue highlight) near the melanocytic gene *GPR143*, and enhancers activated by gained TFE3-peaks (red highlight) near the EMT gene *ZEB1*. Bigwig files of anti-MITF CUT&RUN-seq and ATAC-Seq from *MITF-WT* and *MITF-KO* SKMEL28 cells, as well as ATAC-Seq traces ^4^ from melanocytic (blue), intermediate (black), and mesenchymal-like (red) melanoma cells are shown.

To test the importance of TFE3 in a model MITF-low melanoma cell line, we produced three independent A375 *TFE3-KO* clones (**Figure S6A-C**). Using anti-TFE3 CUT&RUN in A375 cells, we identified 30,342 anti-TFE3 peaks, 66% of which were located within active chromatin (i.e., marked with H3K27Ac). In *TFE3-KO* A375 clones, no anti-TFE3 peaks were present (**Figure 3A,B** and **Figure S6D**), confirming the specificity of the TFE3 antibody. The large majority of active anti-TFE3 peaks in A375 cells coincided with gained-TFE3 peaks in *MITF-KO* cells (e.g., near *ZEB1*) (**Figure 3A,B**). RNA-Seq analysis showed that 3,344 genes had lower expression and 3,206 had higher expression in A375 *TFE3-KO* cells relative to in A375 *TFE3-WT* cells (**Figure 3C, Table S5**). Of those with lower expression in A375 *TFE3-KO* cells, 32% (1023) were flanked by a TFE3 peak (i.e., transcription start site within 100kb), suggesting they are directly activated by TFE3. Gene-set enrichment analysis (GSEA) revealed that this apparently directly-TFE3-activated subset was enriched for genes in the “epithelial mesenchymal transition” (e.g., *MMP3* and *TNFAI3*), “neural crest cell migration in cancer” (e.g. *FSTL3* and *PTHLH*), and “TNF-alpha signaling via NFKB” gene sets (**Figure 3D-F**). In summary, TFE3 activates an invasive gene program in several MITF-low melanoma cell lines as in *MITF-KO* cells.

**Figure 3:**
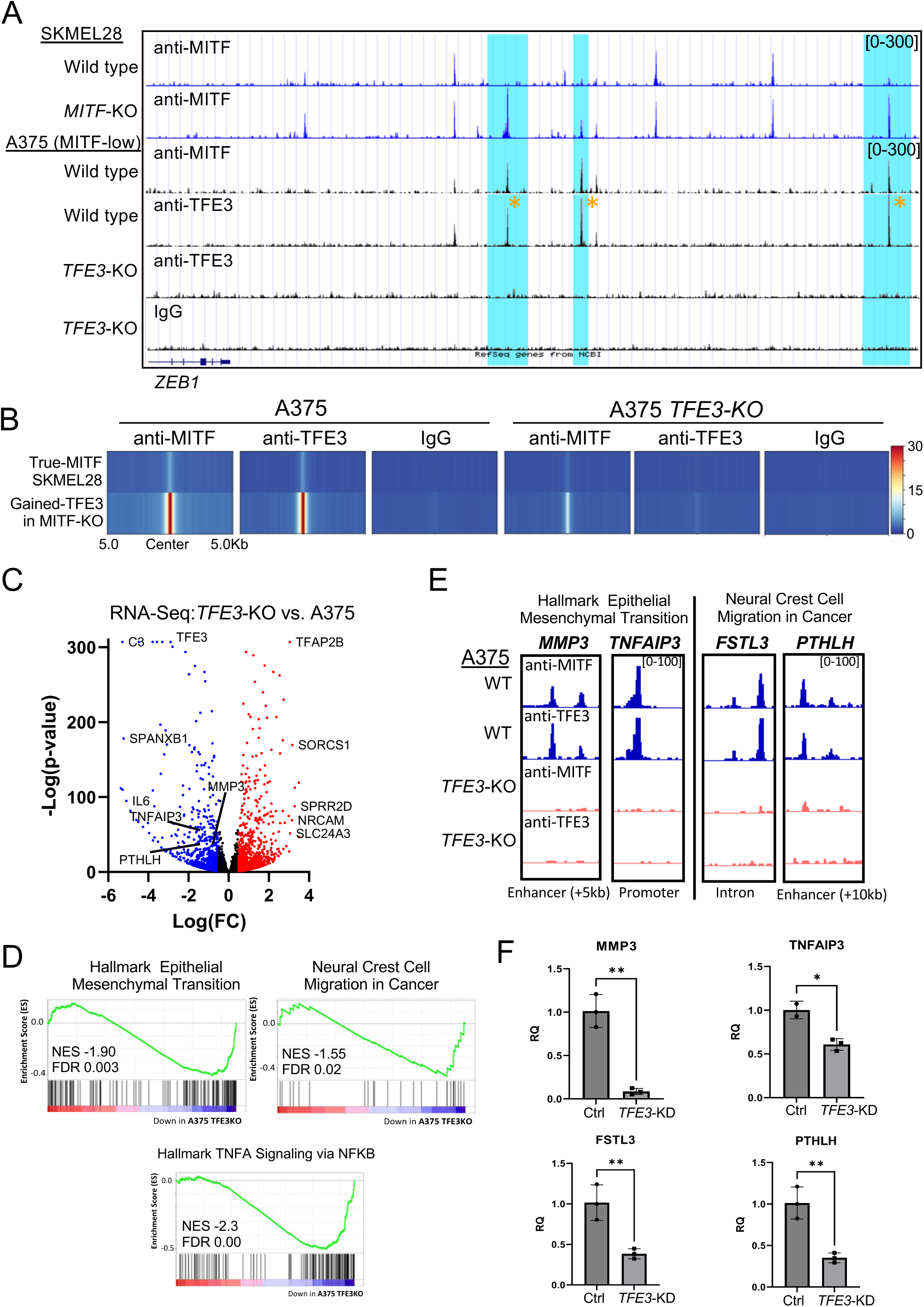
TFE3 establishes a mesenchymal-like transcriptional profile in A375 cell lines. (**A**) Screenshot of IGV genome browser (GRCH37/hg19) visualization of bigwig files generated from anti-MITF CUT&RUN sequencing in *MITF-WT* and *MITF-KO* (D6) SKMEL28 cells, as well as anti-MITF, anti-TFE3, and IgG CUT&RUN sequencing in *TFE3-WT* or *TFE3-KO* A375 cells. Gained paralog peaks in *MITF-KO* cells near the *ZEB1* gene are highlighted in blue, and TFE3 occupancy at gained paralog peaks in A375 cells are indicated by yellow asterisks. (**B**) Density heatmap illustrating anti-MITF, anti-TFE3, and IgG CUT&RUN bigwig signals in *TFE3-WT* or *TFE3-KO* A375 cells, at genomic regions bound by true MITF peaks in *MITF-WT*, and at gained-TFE3 peaks in *MITF-KO* SKMEL28 cells cell lines. (**C**) Volcano plot showing differentially expressed genes (DEGs) with qval <0.05; among these DEGs, 3344 were downregulated (blue) and 3206 upregulated (red) log2-fold change (log2FC) ≥ |1| in *TFE3-KO* vs. *TFE3-WT* A375 cells. (**D**) Enrichment scores for gene ontology (GO) terms and pathways based on gene set enrichment analysis (GSEA) of *TFE3-KO* and *TFE3-WT* A375 cell lines. The enrichment score is indicated by green lines, the number of genes by vertical black lines, and changes in gene expression (positive or negative) by the red to blue heatmap (high to low). (**E**) Screenshot of IGV genome browser (GRCH37/hg19) visualization of bigwig files generated from anti-MITF and anti-TFE3 CUT&RUN sequencing in *TFE3-WT* (blue) and *TFE3-KO* (orange) A375 cells, near the indicated TFE3-activated genes associated with the epithelial to mesenchymal transition (EMT) and neural crest-cell migration in cancer. (**F**) qPCR validation of RNA-Seq data shown in (C). Bar chart representing the relative expression levels of genes identified as directly TFE3-dependent in *TFE3*-WT and *TFE3*-KO A375 cells. Control (Ctrl) samples were transfected with non-targeting gRNAs and CRISPR/Cas9 protein complex, while *TFE3*-KO samples represent bulk CRISPR/Cas9-mediated knockout of TFE3 in A375 cells. Error bars indicate the standard deviation of triplicate experiments. Statistical significance was determined using Student’s t-test: *p < 0.05; **p < 0.01.

### TFE3 promotes invasive behavior of melanoma cell lines and patient-derived xenografts

We next tested whether the invasive phenotype characteristic of MITF-low melanoma depends on TFE3 activity using *in vitro* cell invasion assays. Consistent with predictions based on gene expression profiles ^26^, we observed that in TFE3-high cell lines RPMI- 7951 and A375 the two main isoforms of TFE3 were detected by Western blot. Both the full-length and short-length isoforms of TFE3 were present in cytoplasmic and nuclear fractions (**Figure 4A**). These TFE3-high cells were significantly more invasive than the TFE3-low cell lines SKMEL28 and SKMEL24 (**Figure 4B**). Furthermore, bulk CRISPR/Cas9-mediated knockdown of *TFE3* in RPMI-7951 and A375 cells resulted in a significant reduction in cell invasion (**Figure 4C, Figure S7A-D).** Western blot analysis confirmed the loss of both isoforms of TFE3 (**Figure 4D**). Importantly, KO of *TFE3* in three independent clones of A375 cell lines did not significantly affect cell proliferation relative to in WT A375 cells (**Figure S7E**), suggesting that loss of TFE3 in MITF-low cells impairs migration without affecting cell viability or proliferation *in vitro*.

**Figure 4:**
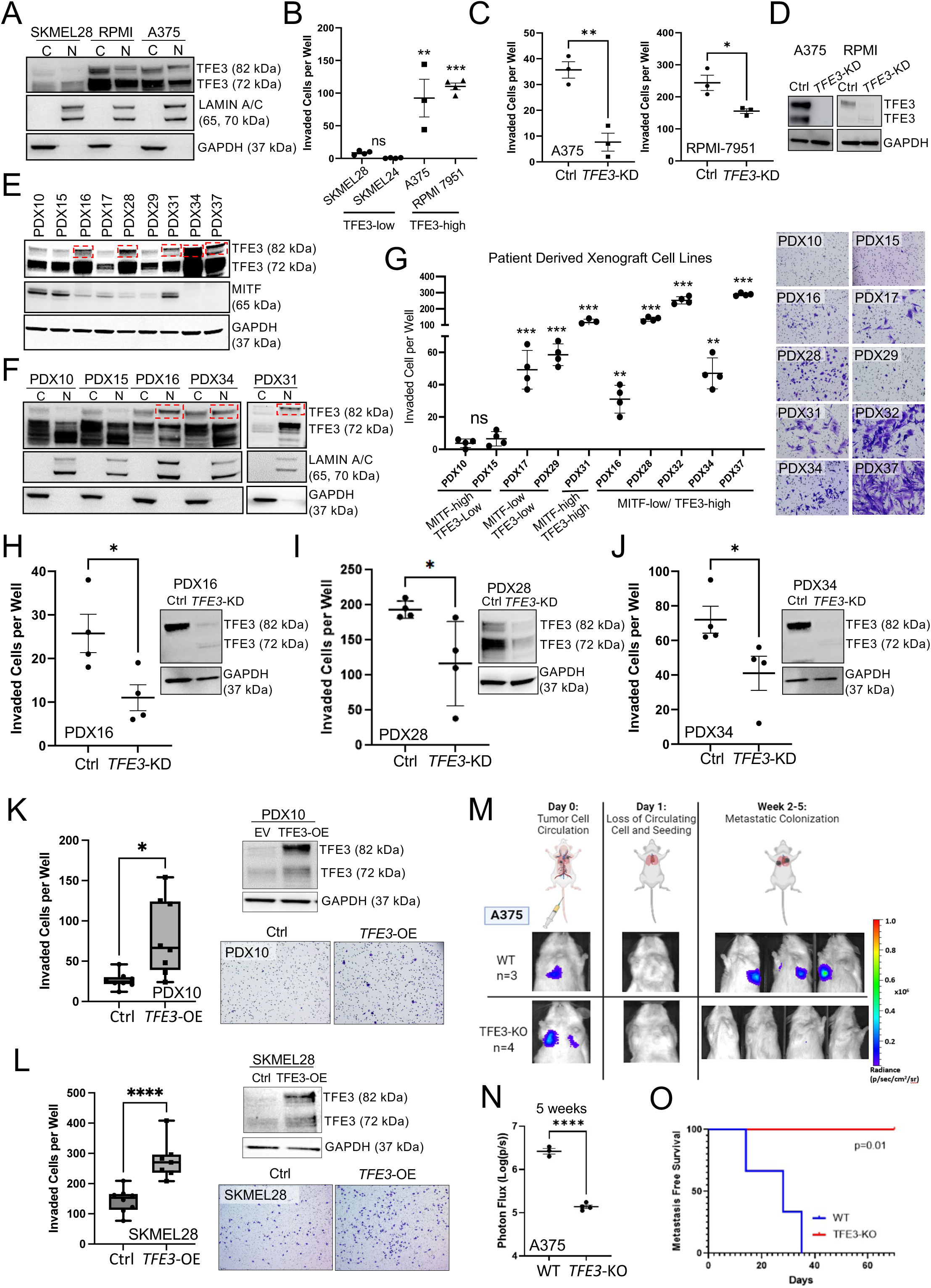
TFE3 accelerates cell invasion and distant lung colonization in MITF-low melanoma cell lines and patient derived xenografts. (**A**) Immunoblotting for anti-TFE3, of cytoplasmic (C) and nuclear (N) fractions of lysates from MITF-high SKMEL24 cells, MITF-low RPMI-7951 and A375 cells. anti-LAMIN A/C and anti-GAPDH served as loading controls for the nuclear and cytoplasmic fractions, respectively. The ∼82 kDa band represents the full-length TFE3 and the ∼72 kDa band represents the short-length TFE3. (**B**) Scatterplot comparing invasive capacity of SKMEL28, SKMEL24, A375 and RPMI-7951cells through Matrigel coated Boydan chambers over 24 hours. Individual dots represent biological experiments with three technical replicates. Statistical analysis was performed using the Student’s t-test. **P-value <0.01, ***P-value <0.001. (**C**) Scatterplot comparing the invasive capacity of A375 and RPMI-7951 cell lines following bulk CRISPR/Cas9-mediated TFE3 knockdown (*TFE3-KD*) or scrambled control (Ctrl), at 24 hours. Individual dots represent biological experiments (n=3) with three technical replicates. Statistical analysis was performed using the Student’s t-test. **P-value <0.01. (**D**) Immunoblotting of anti-TFE3 in A375 and RPMI-7951 cell lines following bulk CRISPR/Cas9-mediated TFE3 knockdown (*TFE3-KD*) or scrambled control (Ctrl). anti-GAPDH served as loading control. (**E**) Immunoblotting of anti-TFE3 and anti-MITF of protein lysates from patient-derived xenograft (PDX) cell lines as labeled, dashed red boxes indicate full-length TFE3. anti-GAPDH served as loading control. (**F**) Immunoblotting of anti-TFE3 from protein lysates of cytoplasmic and nuclear fractions of TFE3-low PDX cell lines (PDX10, PDX15) and of TFE3-high PDX cell lines (PDX16, PDX34 and PDX31). anti-LAMIN A/C and anti-GAPDH served as loading controls for the nuclear and cytoplasmic fractions, respectively. Dashed red boxes indicate full-length TFE3 in nuclear cell fractions. (**G**) Scatterplot comparing invasive capacity of MITF-high and MITF-low PDX cell lines through Matrigel coated Boyden chambers over 24 hours, as labeled. Individual dots represent biological experiments (n=4) with four technical replicates. Statistical analysis was performed using the Student’s t-test. **P-value <0.01, ***P-value <0.001. Representative images of invaded PDX cell lines labeled with crystal violet are shown. (**H-J**) Invasive capacity of (**H**) PDX16, (**I**) PDX28 and (**J**) PDX34, following CRISPR/Cas9-mediated TFE3 knockdown. Control (Ctrl) samples include non-targeting gRNA. Immunoblotting confirms TFE3 knockdown for each cell line. (**K-L**) Boxplot comparing invasive capacity of (**K**) PDX10 and (**L**) SKMEL28 following overexpression of MYC-tagged TFE3 compared to empty vector control (Ctrl). Individual dots represent biological experiments with four technical replicates. Statistical analysis was performed using the Student’s t-test. *P-value <0.05; ****P-value <0.0001. Immunoblotting of anti-TFE3 confirms TFE3 OE for each cell line tested. Representative images of invaded PDX10 and SKMEL28 cell lines labeled with crystal violet as shown. (**M**) Design of lung colonization assay by cells injected via tail vein, with tumor size measured via photon flux (radiance). (**N**) Scatterplot quantitation of size of metastasis in animals shown in (**M**) at 5 weeks post inoculation. Individual dots represent biological replicates. Statistical analysis using the Student’s t-test. **** P-value <0.0001. (**O**) Kaplan Meier curve of metastasis-free survival. Statistical analysis using Log-Rank Test.

We next established 10 patient-derived xenograft (PDX) melanoma cell lines and classified them based on MITF and TFE3 protein expression levels, each either low or high, with TFE3 levels referring to the full-length isoform (**Figure 4E**). Morphologically, the MITF-high PDX cell lines resembled melanocytic melanoma cells, whereas the MITF-low lines displayed an elongated, spindly appearance characteristic of mesenchymal-like melanoma cells (**Figure S8A**), consistent with previous descriptions ^4^. We confirmed that full-length TFE3 localized to the nucleus in three TFE3-high lines (**Figure 4F**). The presence of full-length TFE3 in PDX cell lines significantly correlated with increased cell invasion (r = 0.64, p = 1.40 × 10⁻□ as assessed by Matrigel-coated transwell assay (**Figure 4G**). We further used ANOVA to analyze the relationship between cell invasion and the combined expression of TFE3 and MITF, followed by Tukey’s Honestly Significant Difference test for pairwise comparisons. Cell invasion was significantly higher when both TFE3 and MITF protein expression levels were high compared to conditions where TFE3 expression was low and MITF expression was high (p = 0.034). Additionally, cell invasion was significantly greater when TFE3 expression was high and MITF expression was low compared to conditions where both TFE3 and MITF expression levels were low (p = 0.015).

We next tested necessity and sufficiency of TFE3 for cell invasion *in vitro* by PDX cell lines. Similar to A375 and RPMI-7951, knockdown of *TFE3* by CRISPR/Cas9 in three TFE3-high PDX cell lines, confirmed by Western blot analysis, significantly reduced their cell invasion *in vitro* (**Figure 4H-J**). Conversely, over-expression of full-length, Myc-tagged TFE3 in SKMEL28 and PDX10 cell lines, which otherwise have relatively low invasive behavior (**Figure 4B,G**) and high MITF expression (**Figure 4E, S5E**) significantly increased their invasive behavior (**Figure 4K,L**).

To test the effects of TFE3 on the potential for metastatic colonization we injected *TFE3-WT* and *TFE3-KO* A375 cells labeled with firefly luciferase into the lateral tail vein; from this site, transplanted melanoma cells preferentially colonize the lung ^27^. The time before tumors were detected in the lung and the median time of tumor-free survival were both strongly correlated with genotype. In mice injected with *TFE3-WT* A375 cells, tumors could be detected in the lungs after about 14 days and the median time of tumor free survival was just 5 weeks, while in mice injected with *TFE3-KO* A375 cells tumors were not detected for the duration of the assay (10 weeks) (**Figure 4M-O**).

To evaluate the role of TFE3 in tumor growth *in vivo*, we injected *TFE3*-WT and *TFE3*-KO A375 cells labeled with firefly luciferase into the dermal layer of mouse skin.

Consistent with our *in vitro* assays, where cell growth was not affected by the loss of *TFE3* (**Figure S7E**), tumor growth *in vivo* was comparable between the two genotypes at 4 weeks. However, tumor growth by *TFE3-KO* cells was significantly delayed compared to *TFE3-WT* cells in weeks 1 to 3 (**Figure S9**). In conclusion, TFE3 enhances tumor growth and promotes metastatic colonization of distant organs.

### MITF promotes cytoplasmic retention and lysosome-mediated degradation of TFE3 through activation of FNIP2-mediated non-canonical mTOR signaling

How does loss of MITF lead to increased TFE3 activity? *TFE3* mRNA levels by RNA-Seq were comparable in *MITF-WT* and *MITF-KO* (D2 and D6) SKMEL28 cells arguing against an effect of MITF on *TFE3* transcription or RNA stability. However, as mentioned above, in Western blots we noticed a trend of negatively correlated levels of full-length TFE3 and MITF (**Figure 4A and 4E-F**), with the exception of PDX31, which had high protein expression of both full-length TFE3 and MITF (**Figure 4E-F**). Consistent with this trend, although overall levels of TFE3 protein were comparable in cells of the *MITF-WT* and *MITF-KO* (D2, D6) genotypes, the proportion of full-length to short-length TFE3 was higher in *MITF-KO* cell lines, and similar to that of MITF-low A375 cells (**Figure 5A**); interestingly, the proportion of TFE3 located in the nucleus was also higher in *MITF-KO* cell lines (**Figure 5B**). RT-PCR revealed that there are two *TFE3* RNA isoforms (**Figure 5C**). Sequencing revealed that the two variants differ by 65 base pairs at the 3’ end of exon 2, suggesting they are splice variants, and that the longest open reading frame in each variant correspond to proteins of the 51 and 62 kDa, respectively, with the short-length form missing 105 amino acids from the amino terminus of the full-length form (**Figure 5D**). We presume the 72 kDa (short-length) and 82 kDa (full-length) isoforms correspond to the two splice isoforms, each with post translational modifications.

**Figure 5:**
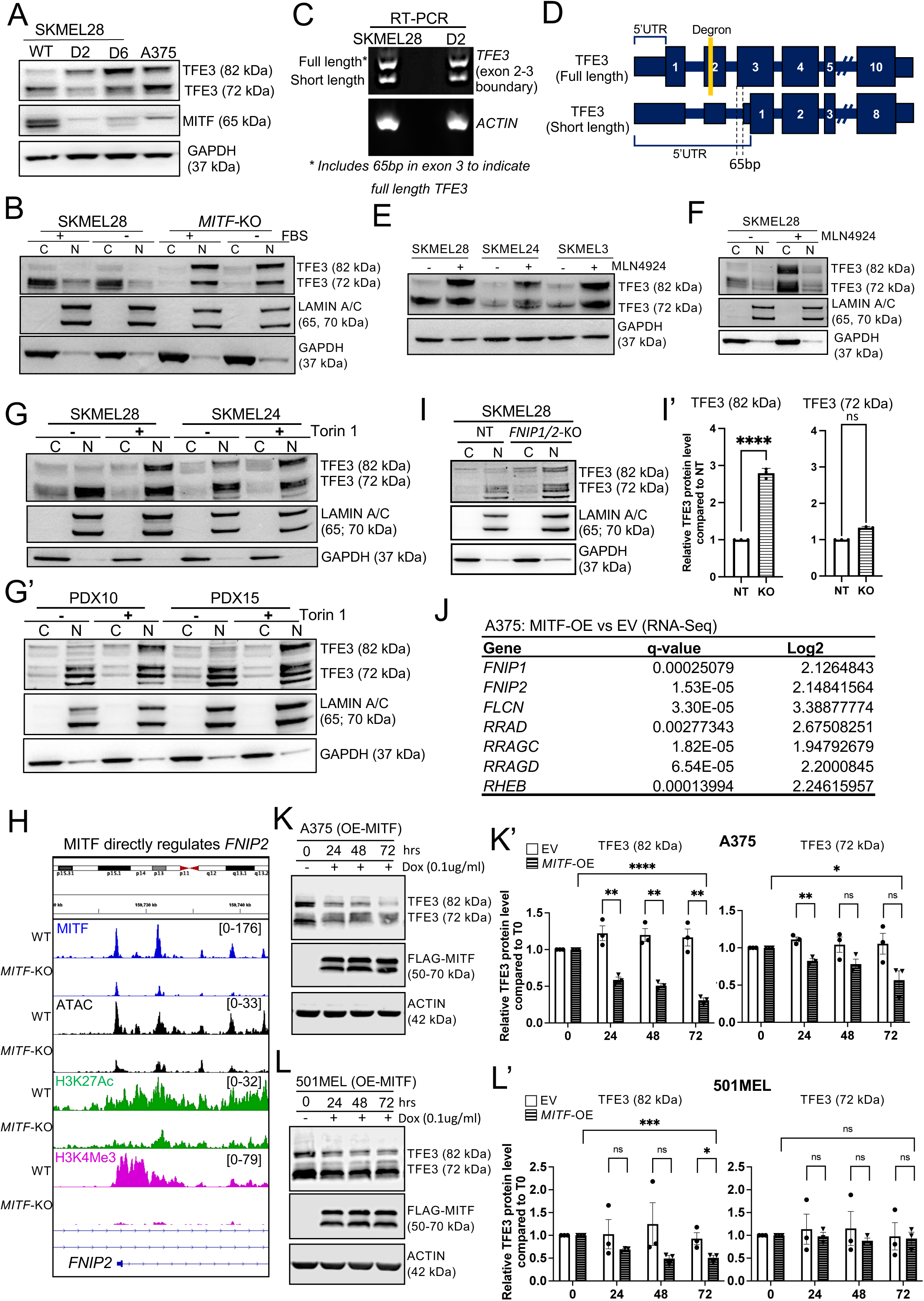
MITF regulates the stability and nuclear localization of full length TFE3 protein by directly activating *FNIP2*-mediated non-canonical mTORC1 signaling. (**A**) Immunoblotting of anti-TFE3 and anti-MITF in lysates from *MITF-WT* and *MITF-KO* (D2 and D6) SKMEL28 cells, and A375 cells. anti-GAPDH served as a loading control. The ∼82 kDa band represents the full-length TFE3 and the ∼72 kDa band represents the short-length TFE3. (**B**) RT-PCR-based visualization of distinct (full-length and short-length) TFE3 transcripts in *MITF-WT* and *MITF-KO* (D2) SKMEL28 cells. Primers targeting actin mRNA were used as a control to ensure equal RNA input and efficient cDNA synthesis across samples. (**C**) Schematic of the full-length and short-length TFE3 transcripts. Indicated are the exons and 5’UTRs, as well as the degron sequence in exon 2 that is unique to the full-length transcript (yellow bar). The TFE3 variants share the same transcriptional start site but have different ribosome entry sites, producing distinct protein products of 72 kDa and 82 kDa. (**D**) Immunoblotting for anti-TFE3, of cytoplasmic (C) and nuclear (N) fractions from lysates of *MITF-WT* and *MITF-KO* (D2) SKMEL28 cells incubated with +/- fetal calf serum (FBS) for 24 hours. LAMIN A/C and GAPDH served as loading controls for the nuclear and cytoplasmic fractions, respectively. (**E**) Immunoblotting for anti-TFE3, in total protein from lysates of SKMEL28, SKMEL24, and SKMEL3 cell lines, -/+ 3 μ MLN4924 for 24 hours. GAPDH served as a loading control. (**F**) Immunoblotting for TFE3, of cytoplasmic and nuclear fractions of lysates from *MITF-WT* SKMEL28 cells following treatment with -/+ 3 μM MLN4924 for 24 hours. LAMIN A/B and GAPDH served as loading controls for nuclear and cytoplasmic fractions, respectively. (**G-G’**) Immunoblotting for anti-TFE3, in cytoplasmic and nuclear fractions of lysates from (**G**) SKMEL28 and SKMEL24 cells, and (**G’**) PDX10 and PDX15 cells, following treatment with Torin 1 (1 uM) for 24 hours. LAMIN A/C and GAPDH served as loading controls for the nuclear and cytoplasmic fractions, respectively. (**H**) Screenshot of IGV genome browser (GRCH37/hg19) visualization of bigwig files generated from anti-MITF, anti-H3K27Ac, and anti-H3K4Me3 CUT&RUN-seq and ATAC-seq, at the *FNIP2* locus in *MITF-WT* or *MITF-KO* (D6) SKMEL28 cells. (**I-I’**) Immunoblotting for anti-TFE3, in cytoplasmic and nuclear fractions of lysates from SKMEL28 following CRISPR/Cas9-mediated knockout of *FNIP1* and *FNIP2*. anti-LAMIN A/C and anti-GAPDH served as loading controls for the nuclear and cytoplasmic fractions, respectively. (**I’**) Histogram showing densitometric analysis of full-length and short-length nuclear TFE3 signals, described in (**I**), quantified using ImageJ. Data represent three independent biological experiments, with individual dots indicating values from each experiment. (**J**) RNA-Seq analysis comparing A375 cells and A375 cells overexpressing MITF. Shown are genes within the mTORC1 pathway that are upregulated upon MITF overexpression, with q-values and Log2 fold change (Log2FC) indicated. RNA-Seq was analyzed from ^22^. (**K-L**) Immunoblotting for anti-TFE3 and anti-MITF in doxycycline-inducible (**K**) A375 cells and **(L**) 501MEL cells at 24, 48, and 72 hours following either mock treatment or doxycycline treatment at the indicated concentrations. (**K’-L’**) Histograms showing densitometric analysis of full-length and short-length TFE3 signals in doxycycline treated (K’) A375 and (L’) 501MEL cell lines, as described in (K) and (L), and normalized to the empty vector controls, across 72 hours. Quantification was performed using ImageJ. Data represent three independent biological experiments (n=3), with individual dots indicating values from each experiment. Statistical analysis using the Student’s t-test. * P-value <0.05; **P-value <0.01

Because the RNA isoforms are expressed at similar ratios in *MITF-WT* and *MITF-KO* SKMEL28 cells (**Figure 5C**) it is unlikely that differential splicing accounts for the different ratios of the protein isoforms in the two genotypes.

In fetal-kidney-derived HEK293 cells, non-canonical mTOR activity phosphorylates full-length TFE3, recruiting Cullin-RING E3 Ubiquitin Ligase 1 (CUL1) to ubiquitylate TFE3, thereby targeting it for lysosomal degradation ^28^. The CUL1-targeted peptide within TFE3, called a degron, is absent from TFE3 translocations present in renal cell carcinoma, accounting for their stable expression and oncogenicity ^28^. This degron is also absent from the short-length isoform of TFE3 (**Figure 5C**), and from the variant of MITF that is present in melanocytes and melanoma cells ^11,28^. Non-canonical mTOR activity also phosphorylates amino-acids present in both isoforms of TFE3 resulting in binding by 14-3-3 and cytoplasmic retention ^29^.

To determine whether TFE3 levels are sensitive to CUL1 activity in melanoma, we inhibited CUL1 in three TFE3-low cell lines by treating them with MLN4924, a selective inhibitor of NEDD8-activating enzyme ^30^. Treatment with MLN4924 led to an increase in levels of the full-length form of TFE3 (**Figure 5E)**. Of note, treatment with MLN4924 did not affect subcellular localization of these forms (**Figure 5F**), suggesting that the stabilized full-length TFE3 protein is bound by 14-3-3 proteins and maintained in the cytoplasm.

Next, to determine whether the mTOR pathway regulates the stability and subcellular distribution of full-length TFE3 in melanoma, we applied the mTOR inhibitor Torin1 or the vehicle DMSO to five MITF-high/TFE3 low cell lines – comprising SKMEL28, SKMEL24, SKMEL3, PDX10 and PDX15. In all cell lines, Torin1 treatment increased the amount of full-length TFE3 and the proportion of it localized to the nucleus; such treatment did not grossly affect the amount of the short-length isoform, nor its cellular distribution, which was largely in the nucleus (**Figure 5G-G’**, bulk data for SKMEL3 in **Figure S10A**). The mTOR pathway responds to cellular nutrition, and in an immortalized human retinal pigmented epithelial cells, serum deprivation for 24 hours induced nuclear localization of TFE3 through inhibition of 14-3-3 binding ^31^. We found that serum starvation of A375 cells (**Figure S10B-C**), but not of the SKMEL28 cells (**Figure 5B**) significantly affected the nuclear localization of TFE3. In summary, the results suggest that in melanoma cells, as in HEK293 cells, mTOR-mediated CUL1 activity destabilizes full-length TFE3, and mTOR activity promotes retention of full-length TFE3 in the cytoplasm.

The results described above led us to assess whether expression of any components non-canonical mTOR activity depend on MITF. Such components include a complex containing folliculin (FLCN) and folliculin-interacting proteins 1 and 2 (FNIP1 and FNIP2) (reviewed in ^32^). In induced pluripotent stem cells this complex promotes phosphorylation and cytoplasmic retention of TFE3 ^25^. Intriguingly, the expression of several members of the FLCN and mTOR pathway is significantly lower in *MITF-KO* (D2 and D6) vs. *MITF-WT* cells (**Table S6**). In particular, an *FNIP2* promoter and several enhancers within the gene body appear to be directly activated by MITF, because they are bound by true-MITF peaks in SKMEL28 cells, and because H3K27Ac and H3K4me3 signals overlying these loci are much lower in *MITF-KO* D6 cells (**Figure 5H**). To test whether the knockout of *FNIP2* is sufficient to stabilize full-length TFE3 protein in high-MITF melanoma, we used CRISPR-Cas9 reagents to knockout *FNIP2* in SKMEL28 cells. Western blot analysis revealed a modest increase in full-length TFE3 in cells transfected with these reagents relative to untransfected cells (**Figures S11A-B**); however, the results were inconsistent. Given that FNIP1 and FNIP2 redundantly activate non-canonical mTORC1 signaling, we designed gRNAs to target both *FNIP* paralogs. Upon depletion of both *FNIP1* and *FNIP2*, full-length TFE3 was consistently stabilized across replicate experiments (**Figures 5I-I’**). Of note, expression of both *YWHAH* (14-3-3 Eta) and *YWHAQ* (14-3-3 Tau), which mediate cytoplasmic localization of many proteins, also appear to be directly activated by MITF in SKMEL28 cells (**Table S6**).

Finally, we tested whether overexpression of MITF in MITF-low/TFE3-high cell lines would result in upregulation of genes encoding members of the mTORC1 pathway and a reduction of TFE3 protein. To address the former we utilized our previously published RNA-Seq data ^22^ were A375 cells were transfected with a plasmid driving constitutive expression of *MITF*. Re-analysis of this dataset revealed that *FNIP2* and mTORC1- related genes (*FNIP1, FLCN, RRAD, RRAGC, RRAGD, RHEB*) were upregulated in these cells relative to in empty-vector-transfected cells (**Figure 5J**). To test whether overexpression of MITF affected TFE3 protein levels, we generated stable transgenic A375 and 501MEL cells with doxycycline-inducible expression of FLAG-tagged MITF (**Figure 5K-L)**. Western blot and densitometry analysis revealed doxycycline treatment had no effect on TFE3 levels in empty-vector transfected control cells but caused a significant reduction in TFE3 levels in transgenic cells, with a stronger effect on the full-length form than on the short-length form (**Figure 5K,K’, 5L,L’** and **Figure S11**).

## Discussion

The data reported herein demonstrate that TFE3 promotes the expression profile and behaviors characteristic of MITF-low invasive melanoma cells, and reveal a mechanism by which MITF normally suppresses these qualities. Our investigation of transcription factor binding profiles and chromatin accessibility in *MITF-WT* and *MITF-KO* melanoma cells showed that in an MITF-high melanoma cell line MITF activates enhancers at genes associated with pigmentation and differentiation, whereas in cells lacking MITF, its paralog TFE3 activates the expression of genes known to promote migration and stem- cell-like features. Further, we found that knockdown or knockout of *TFE3* in aggressive MITF-low cell lines and PDXs suppressed their migration *in vitro* and lung colonization *in vivo.* Loss of TFE3 expression may also have decreased their stem-cell like qualities, as TFE3 directly occupied genes associated with pluripotency (*SOX2*, *KLF4* and *WNT* family members), as it does in naïve embryonic stem cells ^25^. Levels of MITF and TFE3 were previously reported to be negatively correlated ^33^, and we found that both the stability and nuclear localization of TFE3 protein were altered by the knockout of MITF.

We also found that MITF directly activates the expression of both members of the 14-3-3 protein family and of *FNIP2*. The former potentially bind TFE3 and sequesters it in the cytoplasm ^29^, and the latter cooperates with FNIP1 and FLCN to recruit mTORC1 and TFE3 to the lysosome membrane, promoting degradation of the full-length isoform ^28^.

FLCN-mediated degradation of TFE3 also occurs in hESCs ^25^. Collectively, our data show that TFE3 promotes an invasive phenotype, and that MITF, by promoting expression of *FNIP2* and other components of non-canonical mTORC1 signaling, inactivates TFE3.

Our findings support a role for additional MITF paralogs in melanoma biology, as indicated in a recent study ^34^. While most persistent-paralog peaks are reduced in height in *MITF*/*TFE3* double*-KO* cells, they are not absent, which is consistent with MITF paralogs other than TFE3 binding and regulating gene expression. Also, while we have focused on the importance of gained-TFE3 peaks in regulating gene expression in MITF-low cells, at about 27% of persistent-paralog peaks, there is enhancer activity in *MITF-WT* cells but this activity is higher in *MITF-KO* cells. One explanation for this phenomenon is that MITF, possibly as a heterodimer with a paralog, binds this set of enhancers in WT cells but a paralog binds them in *MITF-KO* cells, and this MITF paralog, or heterodimer of paralogs, activates enhancers more avidly than MITF. At the subset of these enhancers where anti-MITF CUT&RUN peaks are present in *MITF/TFE3* double-*KO* cells, the paralog is unlikely to be TFE3.

Finally, although many studies have reported that inhibiting the mTORC1 pathway in preclinical models of cancer is beneficial ^35–40^, our findings suggest that a more nuanced approach may be more successful. mTORC1 is a signaling hub that integrates environmental cues, including levels of nutrients, metabolic intermediates, and growth factors, to regulate growth and metabolism ^41^. However, canonical and non-canonical mTORC1 activities differ in the mechanisms by which substrate is recruited ^32^.

Substrates of canonical mTORC1 activity include p70 ribosomal S6 kinase (S6K) and eukaryotic initiation factor 4E-binding protein 1 (4EBP1), whereas a non-canonical substrate is TFE3 ^32^. Clearance of FLCN suppresses mTORC1-mediated phosphorylation, and thereby inactivation, of TFE3, but it does not suppress the phosphorylation of its canonical substrates ^42^. Multiple studies have shown that the mTOR kinase is active in malignant melanoma ^39^, and that rapamycin, which inhibits this kinase activity, also inhibits melanoma tumor growth ^43–45^. In one recent case, this inhibitory effect on tumor growth was shown to involve autophagy ^44^. While inhibiting the activity of mTORC1 at its canonical targets may be beneficial to the patient, results presented here suggest that mTORC1 activity towards its non-canonical substrate TFE3 suppresses metastatic behavior. These opposite consequences might guide the design of novel drugs targeting specific mTORC1 activities.

## Limitations of this study

Our study reveals that the proliferative state suppresses the invasive one in part through MITF-mediated degradation of TFE3. However, as not all of the genes upregulated in *MITF-KO* cells were downregulated in *MITF/TFE3* double*-KO* cells, there must be other transcription factors driving the invasive state. These may include other MITF paralogs as recently addressed ^34^. Additionally, we have not addressed what triggers the switch to low MITF activity in melanoma. Environmental cues, such as glucose availability ^17,18^, hypoxia ^19,20^, and inflammatory signaling ^18^, suppress MITF expression in melanoma.

How such tumor microenvironmental cues affect activity of TFE3 and other MITF paralogs require further exploration. Additionally, we have not determined whether gained-TFE3 peaks represent binding by homodimers of full-length TFE3, heterodimers of full-length and short-length TFE3, or heterodimers of TFE3 with other MITF paralogs.

## Supporting information

Supplemental files Chang et al

## Resource Availability

### Lead contact

Requests for further information and resources should be directed to and will be fulfilled by the lead contact, Colin Kenny, PhD

### Materials availability

Plasmid constructs and cell lines generated in this study may be obtained by a request to the lead contact.

### Data and materials availability

All processed and raw high-throughput sequencing data from ATAC-seq, RNA-seq and CUT&RUN-seq experiments were uploaded to NCBI GEO. GEO accession numbers GSE273789 (RNA-Seq) and GSE273815 (ATAC-Seq and CUT&RUN-Seq).

**The following datasets were generated:**

| Author(s) | Year | Title | Dataset | Database and Identifier |
| --- | --- | --- | --- | --- |
| Chang et al. | 2024 | Melanoma plasticity is regulated by mTOR-mediated crosstalk between MITF and TFE3 | **ATAC-seq:** SKMEL28 and *MITF*-KO (D6) cell lines | GSE273815 |
|  |  |  | **CUT&RUN-Seq:** anti-H3K27Ac, H3K27Me3 and anti-H3K4Me3 in SKMEL28 and *MITF*-KO (D2 and/or D6). |  |
|  |  |  | anti-MITF and anti-TFE3 in *MITF*-KO (D2 and D6), D2/*TFE3*-KO, A375 and A375 *TFE3*-KO cell lines |  |
|  |  |  | anti-H3K9Ac in A375 cells. |  |
| Chang et | 2024 | Melanoma plasticity is | **RNA-seq:** | GSE273789 |
| al. |  | regulated by mTOR-mediated crosstalk between MITF and TFE3 | A375 and A375 *TFE3*-KO cell lines. |  |
|  |  |  | SKMEL28, *MITF*-KO (D2) and D2/*TFE3*-KO cell lines. |  |

**The following previously published datasets were used:**

| Author(s) | Year | Title | Dataset | Database and Identifier |
| --- | --- | --- | --- | --- |
| Kenny et al. | 2022 | Cut and Run data of MITF in SKMEL28 cells | **CUT&RUN-seq:** anti-MITF, SKMEL28<br>**ATAC-seq:** SKMEL28<br>**RNA-seq:** SKMEL28 | GSE190610 |
| Wouters et al. | 2021 | Single-cell analysis of gene expression variation and phenotype switching in melanoma | **OmniATAC:** MM001, MM011, MM031, MM074, MM074, MM057, MM087, MM047, MM029, MM099. | GSE134432 |
| Dilshat et al. | 2020 | MITF reprograms the extracellular matrix and focal adhesion in melanoma | **RNA-seq:** SKMEL28 and MITF-KO (D6), A375, A375 MITF-overexpression. | GSE163646 |

## Acknowledgements

We gratefully acknowledge Dr. Christine Blaumueller of the Scientific Editing and Research Communication Core at the University of Iowa Carver College of Medicine for critical reading of the manuscript. We also acknowledge Mr. Christopher Franke and Dr. Nicholas Stone at the University of Iowa College of Medicine for bioinformatic support. We acknowledge lab management by Mr. Frank Radella at University of Washington and Mr. Gregory Bonde at University of Iowa. We also acknowledge experimental support by Mr. Dylan Friend and Mr. Isaac O’Toole at University of Iowa.

## Author Contributions

Conceptualization: RAC, CK

Methodology: JC, KCH, MV, JJ, JY, NIB, CM, RN, AKA, HNV, AB, AH.

Investigation/Data Analysis: JC, KCH, MV, JY, AH, MDH, ES, RJW, RAC, CK

Visualization and Data Interpretation: JC, KCH, MV, JY, AH, MDH, ES, RJW, RAC, CK

Funding acquisition: ES, MDH, RAC, CK

Project administration: CK

Supervision: RAC, CK

Writing—original draft: JC, RAC, CK

Writing—review & editing: JC, KCH, RJW, RAC, CK

Approval of Submission: JC, KCH, MV, JY, NIB, AB, AH, MDH, ES, RJW, RAC, CK

Accountable for Contributions: JC, KCH, MV, JY, NIB, AB, AH, MDH, ES, RJW, RAC, CK

## Inclusion and diversity statement

One or more of the authors of this manuscript self-identifies as an underrepresented ethnic minority and gender minority. We support inclusive, diverse and equitable conduct of research.

## STAR Methods

Research specimens and/or clinical data were obtained through the University of Iowa Holden Comprehensive Cancer Center’s ’Melanoma: Skin and Ocular Tissue Repository’ (MAST), an Institutional Review Board – approved biospecimen repository and data registry (IRB 200804792). Informed consent was obtained from all subjects.

### Cell Culture

The SKMEL28 (HTB-72), A375, RPMI-7951 (HTB-66), SKMEL3 (HTB-69), and SKMEL24 (HTB-71) cell lines were purchased from the American Type Culture Collection (ATCC; Manassas, VA, USA) and maintained at 37°C in a 5% CO_2_ atmosphere, in the medium recommended by the manufacturer. The *MITF-KO* D2 and *MITF-KO* D6 cell lines were a gift from the laboratory of Eirikur Steingrimsson (University of Iceland). SKMEL28 was maintained in RPMI 1640 medium (#11875093, ThermoFisher Scientific) supplemented with 10% fetal bovine serum (FBS) and 1% penicillin-streptomycin. A375 cells were maintained in DMEM medium (#11965092, ThermoFisher Scientific) supplemented with 10% FBS and 1% penicillin-streptomycin.

SKMEL3 and SKMEL24 cells were maintained in EMEM medium (30-2003, ATCC) supplemented with 15% FBS and 1% penicillin-streptomycin. RPMI-7951 cells were maintained in EMEM medium (30-2003, ATCC) supplemented with 10% FBS and 1% penicillin-streptomycin. Patient-derived melanoma cell lines were developed directly from a patient biopsy (PDX28, PDX29, PDX32, PDX34, and PDX37) or a first-generation PDX (PDX10, PDX15, PDX16, PDX17, and PDX31), and cultured in RPMI1640 supplemented with 10% FBS, 1%NEAA.

### Generation and culture of PDX cell lines

Patient tumor samples to be used in PDX models were collected and processed by the University of Iowa College of Medicine Tissue Procurement Core under IRB# 200804792. Tissue collection and distribution were performed in accordance with the guidelines of the University of Iowa Institutional Review Board. Briefly, after tumor samples were collected, they were placed in DMEM containing 5% FBS, 1X Pen/Strep, and 1% fungizone. For initiation of PDX tumors, patient tumor samples were processed by mincing until they passed through an 18 G needle easily. The cells were then suspended in a 1:1 mixture of DMEM (GIBCO) and Matrigel (#354230, Corning), and were then subcutaneously injected into 2 or 3 NOD.Cg-Prkdc^scid^Il2rg^tm1Wjl^/SzJ (NSG) (Jackson Laboratory) mice. For the propagation of samples, tumors were dissociated into single-cell suspensions and each mouse was injected subcutaneously with 1x10^6^ cells. Single-cell suspensions were generated by placing the tumor into a sterile Petri dish containing 10mL collagenase IV solution (9 mL HBSS (Ca^2+^- and Mg^2+^-free), 1 mL collagenase IV (Worthington Biochemical Corp.), and 5 mM CaCl_2_). The tumor samples were first minced and incubated at 37°C for 1 hour. Samples were then further digested in 10 mL 0.05% trypsin (at 37°C for 10 minutes), and passed through an 18 G needle 30 times, after which the suspension was filtered using a 70 μ cell strainer.

### Generation of *TFE3-KO* and *FNIP2*-KO cells and validation of mutations using Sanger Sequencing

CRISPR/Cas9 strategies were utilized to generate *TFE3* knockout (KO) A375, RPMI- 7951, *MITF-KO* (D2 and D6) and PDX cell lines, as well as in the *FNIP2* and *FNIP1* genes in SKMEL28 cells. For *TFE3*, a guide RNA (gRNA) was designed to target exon 4 of *TFE3*, which encodes the helix-loop-helix DNA binding domain ^46^ (**Figure S6A-D**). The gRNA used for *TFE3* was: CTTCTCTGAGCTGGACCCGA (Hs.Cas9.TFE3.1.AC Integrated DNA Technologies INC, IDT). For *FNIP2*, a double-guided approach was used, targeting exons 4 and 6 (NM_001323916), with an intermediate segment of approximately 1800 bp. The *FNIP2* guide RNAs used were: AGGTTACTTCCAGTACGACA and AACATGTTGACATCGGAAGC (Hs.Cas9.FNIP2.1.AK and Hs.Cas9.FNIP2.1.AL Integrated DNA Technologies INC, IDT) (**Figure S11A**). A single guide RNA was designed to target exon 5 of *FNIP1*. The *FNIP1* guide RNA used was: AAAGTGTTTACTGCTCGGAC (Integrated DNA Technologies INC, IDT, Hs.Cas9.FNIP1.1.AA).

In each case, the gRNA was conjugated 1:1 with tracerRNA (crRNA) (IDT) by incubation at 95°C for 5 minutes, followed by a period of rest until it reached room temperature.

Cas9 nuclease (#10007807, IDT) was used to form ribonucleoprotein complexes (RNPs). Cells (1 x 10^6^) were electroporated with RNPs in Gene Pulser electroporation buffer (#1652677, Bio-Rad, Hercules, CA) with electroporation enhancer (#1075915, IDT), using 140V and a 2mm pulse gap. A nontargeting negative control gRNA (#1072544, IDT) was used for all CRISPR/Cas9 KOs.

For Sanger sequencing, genomic DNA (CRISPR bulk or clonal isolates) was isolated from the *TFE3-KO* and *FNIP1/2-KO* cell lines using a DNeasy Blood and Tissue Kit (#69504, Qiagen, Hilden, Germany). Regions of interest were amplified using region-specific primers: For *TFE3*, Fwd GCTAGCTCCATGGCTTAGCG, Rev GAGACGCCAACCACAGAGAT. For *FNIP2*: Fwd TCACCGATGACACTAACGAGA, Rev TGCAGCAGACCTAGAGATGG. For *FNIP1*: Fwd GCAGAACAGCGTATACAGTGT, Rev TGAAGAGCACCAACCCTACA. The amplified DNA was run on a 2% agarose gel at 100V for 20 minutes. The bands were cut out of the gel and DNA was purified using the QIAquick Gel Extraction Kit (#28704, Qiagen) and QIAquick PCR Purification Kit (#28104, Qiagen). Purified DNA fragments were cloned into the pcDNA3.1 (+) vector (#V79020, Invitrogen), which was then used to transform 5-alpha competent cells (#C2987H, New England Biolabs, Ipswich, MA). For *TFE3-KO* each condition, 10 colonies were picked, plasmid DNA was isolated and sequenced by Sanger Sequencing. Purified DNA fragments from the *FNIP1/2-KO* were sent directly for Sanger Sequencing.

### RNA isolation, cDNA synthesis, and RT-qPCR

Cells were grown to 70-80% confluency in 6-well culture plates (Corning) and RNA was isolated by direct lysis using the RNeasy Plus kit (#74034, Qiagen). cDNA was synthesized using the High-Capacity cDNA Reverse Transcription Kit (#4368814, Applied Biosystems). qPCR was performed using Taqman qPCR reagents (#4304437 ThermoFisher Scientific,) using the following probes*: MMP3* (Hs00968305_m1, ThermoFisher Scientific), *FSTL3* (Hs00610505_m1, ThermoFisher Scientific), *TNFAIP3* (Hs00234713_m1, ThermoFisher Scientific), *PTHLH* (Hs00174969_m1, ThermoFisher Scientific) and *ZEB1* (Hs01566408_m1, ThermoFisher Scientific). qPCR reactions were performed in triplicate and relative gene expression was calculated using the D-ΔΔCt method ^47^. The expression of target genes was normalized to the expression of the 18S ribosomal subunit (Hs99999901_s1, ThermoFisher).

### Wound scratch assay

6-well plates were seeded at 8 x 10^5^ cells/well and scratches within confluent monolayers were made with a sterile P200 pipet tip in a cross configuration. For each well, four areas in the cardinal directions were chosen for image analysis. Images were recorded at each chosen area using 4X magnification. Wound closure analysis was performed using ImageJ software (National Institutes of Health, https://imagej.net/ij/) and the wound healing plug-in. Area of the wound was quantified at 0-, 24-, and 48-hour time points to assess gap closure.

### Transwell invasion assay

24-well Boyden transwell chambers with a filter pore size of 8 µm (#353097, Corning, Corning, NY) were coated with growth factor-reduced Matrigel matrix (#354230, Corning) diluted 1:10 in serum-free medium. A suspension of 5,000-15,000 cells in appropriate serum-free medium was added to the Matrigel-coated Boyden chamber. The appropriate medium per cell lines containing 10-15% FBS was added to the lower chamber to serve as a chemoattractant. After 24 hours migrated cells were fixed with crystal violet (with 4% PFA). Brightfield images at 4X and10X magnification were acquired using the EVOS Cell Imaging System (ThermoFisher Scientific) and cells were counted using EVOS cell counting software. Experiments were performed in duplicate, triplicate, or quadruplicate, with four technical replicates per experiment. A Student’s t-test was used to compare outcomes across different conditions.

### Blinding for Cell Biological Assays

To ensure blinding, cell lines were numbered by an independent scientist prior to each *in vivo* and *in vitro* experiment. All counting, plating, injecting, imaging, and post-analyses were performed without knowledge of the experimental conditions. Decoding of the numbering was conducted only after the completion of statistical analysis.

### RNA sequencing, differential gene expression analysis, and Gene Set Enrichment Analysis

RNA-sequencing experiments were performed on SKMEL28, *MITF-KO* (D2), 2 clones of double *MITF/TFE3-KO (D2/TFE3-KO)*, A375 and two clones of *TFE3-KO* A375 cell lines (**Tables S4-S5**). All experiments were performed in triplicate. RNA was isolated using the RNeasy Plus Kit (#74034, Qiagen) and submitted to the Novogene Corporation for library construction and sequencing with 75 bp paired-end reads on Novaseq 6000 platform. All raw sequencing data was assessed by FastQC for quality control. Reads were mapped to the human reference genome hg38 using RNAStar with default parameter settings. The FeatureCounts program was used to calculate read counts for each individual gene from STAR output bam files. Differential gene expression analysis was performed using DEseq2. The adjusted p-values for multiple testing were calculated using the Benjamini-Hochberg (BH) procedure, as implemented in the DESeq2 software package. This method controls the false discovery rate (FDR). Genes with an adjusted p-value < 0.05 were considered statistically significant. To identify enriched gene ontology (GO) terms, differential gene expression between SKMEL28 and D2, and between D2 and D2/TFE3-KO cells were analyzed using GO enrichment analysis (Global Core Biodata Resource). GO terms with FDR values of ≤ 0.05 were considered statistically significant. Additionally, differential gene expression between A375 and A375 *TFE3-KO* clones were analyzed using Gene Set Enrichment Analysis (GSEA) software. GSEA was performed using the signatures “Hallmark Epithelial Mesenchymal Transition”, “Neural Crest Cell Migration in Cancer”, and “Hallmark TNFA Signaling via NFKB” in MSigDB (v2023.1). FDR values of ≤ 0.05 was used for the statistical significance of the enrichment score for each signature.

### CUT&RUN

To identify direct target genes of MITF and TFE3 and histone 3 modifications, we performed CUT&RUN sequencing in SKMEL28, *MITF*-KO (D2), 2 clones of double *MITF/TFE3*-KO (D2/TFE3-KO), A375 and two clones of *TFE3*-KO A375 cell lines, as previously described ^23,48,49^. Log-phase cultures of cells (60-80% confluent) were harvested by cell scraping, centrifuged at 300 g and washed twice in Ca^2+^-free wash-buffer (20 mM HEPES, pH7.5, 150 mM NaCl, 0.5 mM spermidine, and protease inhibitor cocktail, cOmplete Mini, EDTA-free Roche). Cells were counted and 500,000 cells per antibody per condition were used for CUT&RUN experiments (performed in biological duplicates). Preactivated concanavalin A-coated magnetic beads (Bangs Laboratories, Inc) were added to the cell suspension, followed by incubation at 4°C for 10 minutes.

Antibody buffer (wash buffer with 2mM EDTA and 0.025% digitonin) containing anti-MITF (HPA003259, Sigma), anti-TFE3 (#14779, Cell Signaling Technologies (CST)), anti- H3K27Ac (#8173, CST), anti-H3K27Me3 (#9733, CST), anti-H3K4Me3 (#9751, CST), anti-H3K9Ac (#9649, CST), or anti-Rabbit IgG (#12-370, Millipore) was added and cells were incubated overnight at 4°C on a Nutator mixer. The following day, cells were washed in dig-wash buffer (wash buffer containing 0.025% digitonin). pAG-MNase reactions were quenched using 2X Stop buffer (340 mM NaCl, 20 mM EDTA, 4 mM EGTA, 0.05% Digitonin, 100 μ/mL RNAse A, 50 μ /mL Glycogen, and 2 pg/mL sonicated yeast spike-in control). Released DNA fragments were treated with Proteinase K (#19131, Qiagen) for 1 hour at 50°C and purified by phenol/chloroform-extraction and ethanol-precipitated. CUT&RUN experiments were performed in parallel, as positive control and fragment sizes were analyzed using a Qubit Fluorometer and a 2100 Bioanalyzer (Agilent).

### Assay for transposase-accessible chromatin with sequencing (ATAC-Seq)

ATAC-seq was performed according to ^23^ with minor alterations. Briefly, 70,000 *MITF-WT* SKMEL28 and D6 (four replicates) were lysed in ice-cold lysis buffer (10 mM Tris-HCl, pH 7.4, 10 mM NaCl, 3 mM MgCl2, 0.1% NP-40: Sigma). Transposition was performed directly on nuclei using 25μl tagmentation reaction mix (Tagment DNA Buffer #15027866 and Tagment DNA Enzyme #15027865 from the Illumina Tagment DNA kit #20034210). Tagged DNA was subjected to PCR amplification and library indexing, using the NEBNext High-Fidelity 2X PCR Master Mix (#M0451S, New England Biolabs) with Nextera DNA CD Indexes (#20015882, Illumina), according to the following program: 72°C for 5 minutes; 98°C for 30 seconds, 12 cycles of 98°C for 10 seconds, 63°C for 30 seconds, and 72°C for 1 minute. The PCR product was purified with 1.8 times the volume of Ampure XP beads (#A63881, Beckman Coulter). Library quality was assessed using a BioAnalyzer 2100 High Sensitivity DNA Chip (Agilent Technologies).

All DNA libraries that exhibited a nucleosome pattern were pooled and processed for 150 bp paired-end sequencing. GV (Integrated Genomic Viewer, https://igv.org/) was used visualize overlapping regions of CUT&RUN and ATAC-seq data.

### Library preparation and data analysis

CUT&RUN libraries were prepared using the KAPA Hyper Prep Kit (#07962363001, Roche). Quality control post-library amplification was conducted using a Qubit Fluorometer and a 2100 Bioanalyzer for fragment analysis. Libraries were pooled to equimolar concentrations and sequenced with paired-end 150 bp reads on a NovoSeq HiSeq X instrument. Paired-end FastQ files were processed for quality control using the FastQC tool. Reads were trimmed using Trim Galore Version 0.6.3 (developed by Felix Krueger at the Babraham Institute), and Bowtie2 version 2.1.0 ^50^ was used to map the reads against the hg19 genome assembly. The mapping parameters were performed as previously described ^49^. Peak calling was performed with MACS2^51^ and set to narrow peaks for transcription factors and broad peaks for histone modifications. Peaks with FDR values <0.05 were considered statistically significant. Peak intersections were performed with the intersect tool in Galaxy, a minimum of 1bp overlap between peaks was considered an overlapping peak. To visualize signal density across genomic regions, we employed Deeptools^52^. BigWig files were first generated from sorted and indexed BAM files using bamCoverage, normalizing by reads per kilobase per million (RPKM). Signal coverage was then computed using computeMatrix, where genomic regions of interest were defined in a BED file. For transcription start site (TSS)-centered analyses, signal was extracted ±5 kb from the TSS. The resulting matrix was used to generate heatmaps or signal plots with plotHeatmap or plotProfile, which visualized the distribution of signal across the selected regions. For loci-specific visualization, BigWig files were loaded into Integrated Genomics Viewer (IGV) ^53^.

### Cytoplasmic and nuclear fractionation

Cytoplasmic and nuclear protein fractionation and extraction of cell lines were performed using NE-PER Nuclear and Cytoplasmic Extraction Reagents (Cat# 78835. ThermoFisher Scientific). Protein concentration was evaluated with the Pierce BCA Protein Assay Kit (#23225, Invitrogen). Cytoplasmic and nuclear protein components were subsequently run on 4-12% Bis-Tris protein gels (NP0335BOX, Invitrogen), transferred to a PVDF membrane (IB24001 and IB24002, Invitrogen),and incubated with appropriate primary and secondary antibody for the analysis described below. Anti-GAPDH (sc-47724, Santa Cruz) was utilized as cytoplasmic protein loading control and anti-LAMIN A/C (#4777, Cell Signaling Technologies) was used as nuclear protein loading control.

### Drug inhibition in cell lines and PDXs

Cell lines and PDXs were seeded into a 6-well plate (3-5 x 10^5^ cells/well). The following day, cells were treated for 24 hours with cell specific media containing a drug inhibitor. For mTOR inhibition, cells were treated with 1 uM Torin 1 (#4247, Tocris Bioscience).

For CUL inhibition, cells were treated with 3 µM MLN4924 (S7109, Selleck.com). DMSO was added to the media as a control for these experiments. After treatment, both bulk protein and cytoplasmic/nuclear protein were harvested from the cells. Cellular fractionation was performed using the NE-PER Nuclear and Cytoplasmic Extraction Reagents (#78835. ThermoFisher Scientific) and bulk protein was harvested using RIPA

Lysis Buffer (R0278, Sigma-Aldrich) with Halt Protease and Phosphatase Inhibitor Cocktail (#78440, ThermoFisher Scientific). Protein concentration was assessed with the Pierce BCA Protein Assay Kit (#23225, Invitrogen) and were evaluated via Western blot following the techniques outlined in the Immunoblotting section.

### Doxycycline inducible expression of MITF

Doxycycline-inducible A375 and 501MEL cells were seed at a density of 8 × 10□ cells and then treated with 0.1 μg /ml doxycycline to induce expression of the MITF-WT fusion protein tagged with 3XFlag-HA at the C-terminus (MITF-WT-3XFlag-HA) for 24, 48, or 72 hours before harvesting. Cells were lysed in SDS sample buffer (2% SDS, 5% 2- mercaptoethanol, 10% glycerol, 63 mM Tris-HCl, 0.0025% bromophenol blue, pH 6.8). Western blot analyzed protein expression using FLAG antibodies (#F3165, Sigma). Actin was a loading control detected with anti-β-actin antibodies (#4970, Cell Signaling). Proteins were separated via SDS-PAGE and transferred onto 0.2 μ PVDF membranes (#88520, ThermoScientific). The membranes were blocked in T-TBS (20 mM Tris, pH 7.4; 150 mM NaCl; 0.01% Tween 20) containing 5% BSA and probed with specific primary antibodies. After three 10-minute washes with T-TBS, membranes were incubated for 1 hour at room temperature with DyLight 800 anti-mouse (#5257, Cell Signaling Technologies (CST)) or DyLight 580 anti-rabbit IgG (#5366, CST) secondary antibodies. Protein bands were visualized using the Odyssey CLx Imager (LI-COR Biosciences) and quantified with Image Studio (version 2.0) and ImageJ software (National Institutes of Health, https://imagej.net/ij/).

### Immunoblotting

Cells were resuspended in RIPA Lysis Buffer (R0278, Sigma-Aldrich) with Halt Protease and Phosphatase Inhibitor Cocktail (#78440, ThermoFisher Scientific). Protein concentration was assessed using the Pierce BCA Protein Assay Kit (#23225, Invitrogen). NuPAGE LDS Sample Buffer and Reducing Buffer (NP0007 & NP0009, Invitrogen) were added to equal amounts of protein (15-30 mg) and samples were boiled at 95°C for 5 minutes. Protein samples were loaded on 4-12% Bis-Tris gels (NP0335BOX, Invitrogen) and then transferred to polyvinylidene fluoride (PVDF) membranes (IB24001 and IB24002, Invitrogen), which were incubated with primary antibodies overnight at 4°C. The membranes were washed three times with TBS-T and then incubated with horseradish peroxidase-conjugated anti-rabbit or anti-mouse (#7074 & #7076, Cell Signaling Technology). Signal was then developed using SuperSignal West Femto or West Dura (#34095 or #34075, Thermo Scientific). The following antibodies were used for Western blotting:

### Cell Proliferation Assay (MTT)

Cells were seeded at 1,000 cells per well in 96-well plates, in a set of 6 technical replicates. Every 24 hours, cells were incubated in fresh medium with 0.5 mg/ml MTT reagent (#M6494, ThermoFisher) for 2 hours at 37°C and 95% CO_2_, and then solubilized for 10 minutes in 200 ul DMSO. Measurements were obtained on an Infinity M Plex (Tecan: Switzerland) plate reader, at an absorbance of 540 nm. Measurements were taken daily for one week and normalized to absorbance measurements taken at Day 0.

### Intradermal and tail vein injection

A375 cell lines with stable luciferase expression were grown in culture. To assess tumor growth in vivo following deletion of TFE3, 2 x 10^6^ A375 Wild Type and *TFE3-KO* cells were harvested for each injection at subconfluent conditions. Prior to intradermal injection, the backs of NOD/SCID mice were shaved and sterilely prepped. Using a 25- gauge needle, cells suspended in Matrigel with Growth Factor (#354234, Corning) were injected into the intradermal space on the back of each mouse. These mice were inspected daily for tumorigenesis and growth and were imaged using an In Vivo Imaging System (IVIS) for small animals weekly for 4 weeks. After 4 weeks, all mice were euthanized. To assess metastatic colonization, 1 x 10^6^ cells were harvested and injected into the lateral tail vein of immunocompromised NOD/SCID mice. The mice were imaged using an *in vivo* imaging system (IVIS) for small animals on Day 0, to visualize circulating tumor cells and confirm that injection was accurate. Mice were also imaged on Day 1 to confirm that signal was lost, and then weekly to assess metastatic colonization and metastatic growth. To assess tumor growth *in vivo* following deletion of TFE3, A375 and *TFE3-KO* cells (2 x 10^5^) were harvested and injected intradermally into NOD/SCID mice.

Mice were imaged over three weeks to assess tumorigenesis and growth. Mice were euthanized at 4 weeks and imaged using the IVIS system. All mouse experiments were approved by and performed in accordance with the University of Iowa IACUC.

### Electroporation for dual luciferase reporter assay and plasmid overexpression

Each firefly reporter construct for SERPIN3A was co-transfected with a constitutively driven Renilla luciferase plasmid for dual luciferase assays. Briefly, SKMEL28, *MITF*-KO (D2) and double D2/*TFE3*-KO cells were electroporated using the BTX ECM 830 Square Wave Electroporation System (BTX, Harvard Bioscience) and incubated for 36-48 hours. Cells were washed with DPBS (#14190144, Gibco) and media was changed 24 hours after electroporation. We used a dual-luciferase reporter assay system (Promega, Catalog no. E1910) and FB12 Luminometer (Berthold Detection Systems) to evaluate the luciferase activity following the manufacturer’s instructions. Relative luciferase activity was calculated as the normalized values of the Firefly to the Renilla enzyme activities. Three independent measurements were performed for each transfection group, and 2-3 biological replicates were performed. All results were presented as mean ± Standard deviation (s.d.). Statistical significance was determined using the Student’s t test (two-tailed). To overexpress TFE3 in SKMEL28 and PDX10, 4 x 10^5^ cells were transfected with either TFE3-MYC plasmid or piggyBAC-CMV-GFP. Cells were washed with DPBS buffer (#14190144, Gibco), resuspended in 400 uL of ice-cold Gene Pulser electroporation buffer (#1652677, Bio-Rad, Hercules, CA) and mixed with 20 ug of plasmid. The cell and plasmid mixture was added to a 2 mm cuvette (BTX420, BTX) and electroporated using the BTX ECM 830 Square Wave Electroporation System (BTX, Harvard Bioscience). The electroporation settings were as follows:

|  |  |
| --- | --- |
| Voltage | 85 V |
| Duration | 70 msec |
| Number of pulses | 1 |
| Pulse interval | 0.0 sec |
| Electrode gap | 2.0 mm |

Cells were plated into a 6-well plate and incubated under standard cell culture conditions for 48 hours. Cells were washed with DPBS (#14190144, Gibco) and fresh media was added 24 hours post-electroporation.

### Statistics and reproducibility

Statistical analysis was performed using GraphPad Prism 9 statistical software (La Jolla, CA). Tumor-free survival (TFS) was calculated using the Kaplan-Meier method and significance was evaluated by the log rank test. Statistical significance was determined using several statistical tests such as hypergeometric tests, Fisher-exact tests, Mann-whitney U test and ANOVA, as appropriate, with all statistical tests using two sided tests with p < 0.05 considered statistically significant. Biological analyses was performed on a minimum of triplicate samples to ensure reproducibility.

## Supplemental Tables

**Table S1:** Differential gene expression and anti-MITF and anti-TFE3 target genes in SKMEL28 and *MITF*-KO (D2, D6) cell lines.

**Table S2:** Differential gene expression comparing double *MITF/TFE3*-KO (clone 13) and *MITF*-KO (D2) cell lines.

**Table S3:** Differential gene expression comparing double *MITF/TFE3*-KO (clone 3) and *MITF*-KO (D2) cell lines.

**Table S4:** Upregulated genes in *MITF*-KO cells vs. SKMEL28 cells that are associated with naïve human embryonic stem cell identity.

**Table S5:** Differential gene expression comparing *TFE3*-KO and A375 cell lines.

**Table S6:** Downregulated genes in *MITF*-KO cells vs. SKMEL28 cells within the mTORC1 signaling pathway.

## Supplemental Figures

**Figure S1: Multiple sequence alignment of members of the MIT/TFE transcription factor family and the anti-MITF antibody immunogen sequence**.

**Figure S2: Overlap of anti-MITF CUT&RUN peaks in D2 and D6 cell lines.**

**Figure S3: Characterization of anti-MITF CUT&RUN peaks in melanoma cells.**

**Figure S4: Cross-reactivity of anti-MITF antibodies with TFE3.**

**Figure S5: TFE3 is a transcriptional activator in *MITF-KO* (D2 and D6) cell lines.**

**Figure S6: Isolation of *TFE3-KO* clonal cell lines and anti-MITF CUT&RUN-seq.**

**Figure S7: Cell biological endpoint assays in *TFE3-KO* cells**

**Figure S8: Brightfield images of PDX cell lines.**

**Figure S9: Imaging of tumor xenograft injections at 1 week and 3 weeks post injection.**

**Figure S10: TFE3 response to serum depletion in A375 cell lines.**

**Figure S11: Knockout of FNIP2 from MITF-high SKMEL28 cell lines.**

**Figure S12: Overexpression of MITF in A375 and 501MEL cell lines.**

