## Supplemental files Chang et al for "Antagonistic Roles for MITF and TFE3 in Melanoma Plasticity"

```
MITF   Query_10001  1    [ 51]MTSRILLRQQLMREQMQEQERREQQQKLQAAQFMQqrvPVSQTPAINV   SVPTTLPSATQVPMEVLKVQTHLE  123
TFE3   Query_10002  1    [109]SSSRVLLRQQLMRAQAQEQERRERREQAAAAPFPS---PAPASPAISV[ 6]GHTLSRPPPAQVPREVLKVQTHLE  184
TFEB   Query_10003  1         MASRIGLRMQLMREQAQQEEQRERMQQQAVMHYMQqq-QQQQQQQLGG[ 6]NTPVHFQSPPPVPGEVLKVQSYLE  77
TFEC   Query_10004            ------------------------------------------------    ------------------------
Sigma  Query_10005            ------------------------------------------------    ------------------------

       Query_10001  124  NPTKYHIQQAQRQQVKQYLSTTLANKHANQVLSLPCPNQPGDHVMPPVPG------------SSAPNSPMAMLTLNSNCE  191
       Query_10002  185  NPTRYHLQQARRQQVKQYLSTTLGPKLASQALTPP----PGPASAQPLPAPEAAHTT--GPtGSAPNSPMALLTIGSSSE  258
       Query_10003  78   NPTSYHLQQSQHQKVREYLSETYGNKFAAHISPAQGSPKPPPAASPGVRAGHVLSSS--AG-NSAPNSPMAMLHIGSNPE  154
       Query_10004  1    -------------------------MTLDHQIINPTLKWSQPAVPSGGPLVQHAHTTldSDaGLTENPLTKLLAIGKEDD  55
       Query_10005       --------------------------------------------------------------------------------

       Query_10001  192  KE[28]MDDVIDDIISLESSYNEEILGLMD---PALQMANTLPVSGNLIDLYGNQGLPPPGLT--ISNSCPANLPNiKRE  290
       Query_10002  259  KE    IDDVIDEIISLESSYNDEMLSYLPggtTGLQLPSTLPVSGNLLDVYSSQGVATPAIT--VSNSCPAELPNiKRE  332
       Query_10003  155  RE    LDDVIDNIMRL-----DDVLGYIN---PEMQMPNTLPLSSSHLNVYSSDPQVTASLVgvTSSSCPADLTQ-KRE  221
       Query_10004  56   NA[ 3]MEDVIEDIIGMESSFKEE------gadSPLLMQRTL--SGSILDVYSGEQGISPINMglTSASCPSSLPM-KRE  125
       Query_10005       --    --------------------------------------------------------------------------

       Query_10001  291  LT[6]ESEARALAKERQKKDNHNLIERRRRFNINDRIKELGTLIPKSNDPDMRWNKGTILKASVDYIRKLQREQQRAKEL  373
       Query_10002  333  IS   ETEAKALLKERQKKDNHNLIERRRRFNINDRIKELGTLIPKSSDPEMRWNKGTILKASVDYIRKLQKEQQRSKDL  409
       Query_10003  222  LT   DAESRALAKERQKKDNHNLIERRRRFNINDRIKELGMLIPKANDLDVRWNKGTILKASVDYIRRMQKDLQKSREL  298
       Query_10004  126  IT   ETDTRALAKERQKKDNHNLIERRRRYNINYRIKELGTLIPKSNDPDMRWNKGTILKASVEYIKWLQKEQQRAREL  202
       Query_10005       --   ---------------------------------------------------------------------------

MITF   Query_10001  374  ENRQKKLEHANRHLLLRIQELEMQARAHGLSLIPSTGLCSPDLVNRIIKQE-PVLENCSQDL     LQHHADLTCTTTLD  448
TFE3   Query_10002  410  ESRQRSLEQANRSLQLRIQELELQAQIHGLPVPPTPGLLSL------------ATTSASDSL     KPEQLDIE----EE  469
TFEB   Query_10003  299  ENHSRRLEMTNKQLWLRIQELEMQARVHGLPTTSPSGMNMAELAQQVVKQElPSEEGPGEAL[14]LPPQAPLPLPTQPP  388
TFEC   Query_10004  203  EHRQKKLEQANRRLLLRIQELEIQARTHGLPTLASLG--TVDLGAHVTKQQsHPEQNSVDYC     --QQLTVSQGPSPE  274
Sigma  Query_10005  1    ------------HLLLRIQELEMQARAHGLSLIPSTGLCSPDLVNRIIKQE-PVLENCSQDL     LQHHADLTCTTTLD  63

MITF   Query_10001  449  LTDGTITFNNNLGTGTEANQA---YSVPTKMGSK-     LEDILMDD     TLSPV-GVTDPLLSSVSPGASKTSSRRSS  515
TFE3   Query_10002  470  GRPGAATFH--VGGGPAQNAP---HQQPPAPPSDA[21]LEDILMEE[11]ALSPLrAASDPLLSSVSPAVSKASSRRSS  568
TFEB   Query_10003  389  SPFHHLDFSHSLSFGGREDEGppgYPEPLAPGHGS[ 9]LDLMLLDD     SLLPL--ASDPLLSTMSPEASKASSRRSS  467
TFEC   Query_10004  275  LCDQAIAFSDPLSYFTDLSFS-------AALKEEQ[ 1]LDGMLLDD     TISPF--GTDPLLSATSPAVSKESSRRSS  338
Sigma  Query_10005  64   LTDGTITFNNNLGTGTEANQA---YSVPTKMGSK-     LEDILMDD     TLSPV-GVTDPLLSSVSPGASKTSSRRSS  130

MITF   Query_10001  516  MSMEETEHTc  525
TFE3   Query_10002  569  FSMEEES---  575
TFEB   Query_10003  468  FSMEEGDVL-  476
TFEC   Query_10004  339  FSSDDGDEL-  347
Sigma  Query_10005  131  MSMEETEHT-  139
```

**Figure S1**

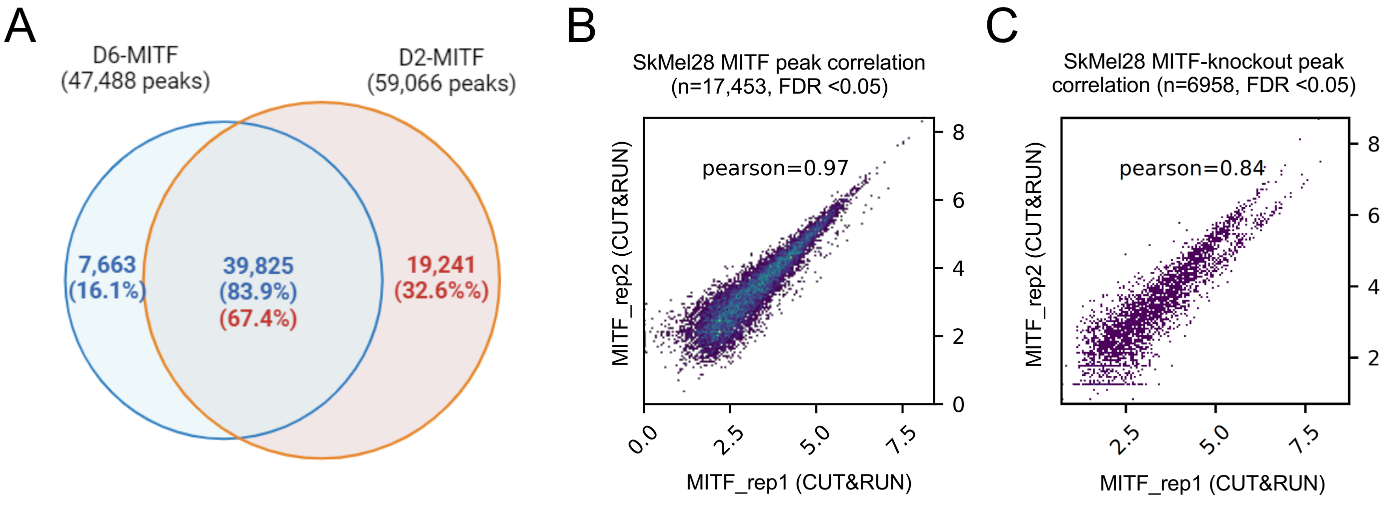
A
D6-MITF
(47,488 peaks)
D2-MITF
(59,066 peaks)
7,663
(16.1%)
39,825
(83.9%)
(67.4%)
19,241
(32.6%%)
B
SkMel28 MITF peak correlation
(n=17,453, FDR <0.05)
pearson=0.97
MITF_rep2 (CUT&RUN)
MITF_rep1 (CUT&RUN)
C
SkMel28 MITF-knockout peak
correlation (n=6958, FDR <0.05)
pearson=0.84
MITF_rep2 (CUT&RUN)
MITF_rep1 (CUT&RUN)

**Figure S2**

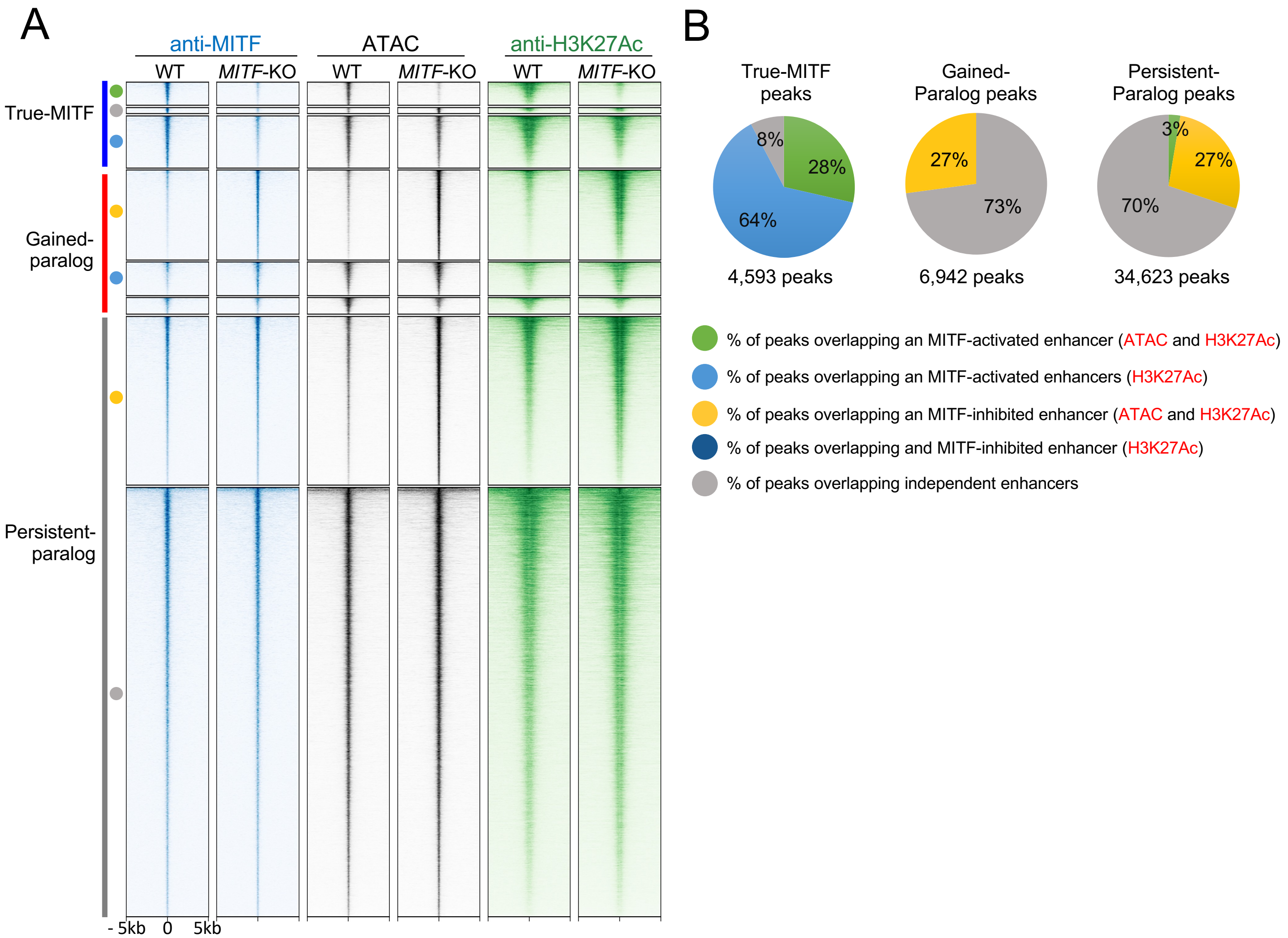


C

True MITF peaks
Gene Ontology analysis (Panther)

| GO | No. of genes | FDR |
|---|---|---|
| Developmental processes | 1103 | $2.6x10^{-15}$ |
| Cell differentiation | 710 | $1.7x10^{-09}$ |
| Developmental pigmentation | 21 | $3.7x10^{-05}$ |
| Regulation of cytokine production | 173 | $8.9x10^{-05}$ |
| Melanocyte differentiation | 10 | $2.6x10^{-03}$ |
| Organelle localization | 119 | $5.0x10^{-03}$ |
| Pigment cell differentiation | 14 | $8.4x10^{-03}$ |
| Melanin biosynthetic process from tyrosine | 4 | $1.9x10^{-02}$ |

D

Gained-paralog peaks
Gene Ontology analysis (Panther)

| GO | No. of genes | FDR |
|---|---|---|
| Regulation of cell migration | 266 | $1.5x10^{-17}$ |
| Extracellular matrix organization | 87 | $1.0x10^{-13}$ |
| Regulation of metabolic process | 1287 | $2.4x10^{-13}$ |
| Regulation of metabolic process | 1287 | $2.4x10^{-13}$ |
| Kidney development | 97 | $8.9x10^{-09}$ |
| Regulation of MAPK cascade | 170 | $4.5x10^{-08}$ |
| Regulation of epithelial to mesenchymal transition | 35 | $3.8x10^{-4}$ |
| Wound healing | 78 | $4.0x10^{-03}$ |

Figure S3

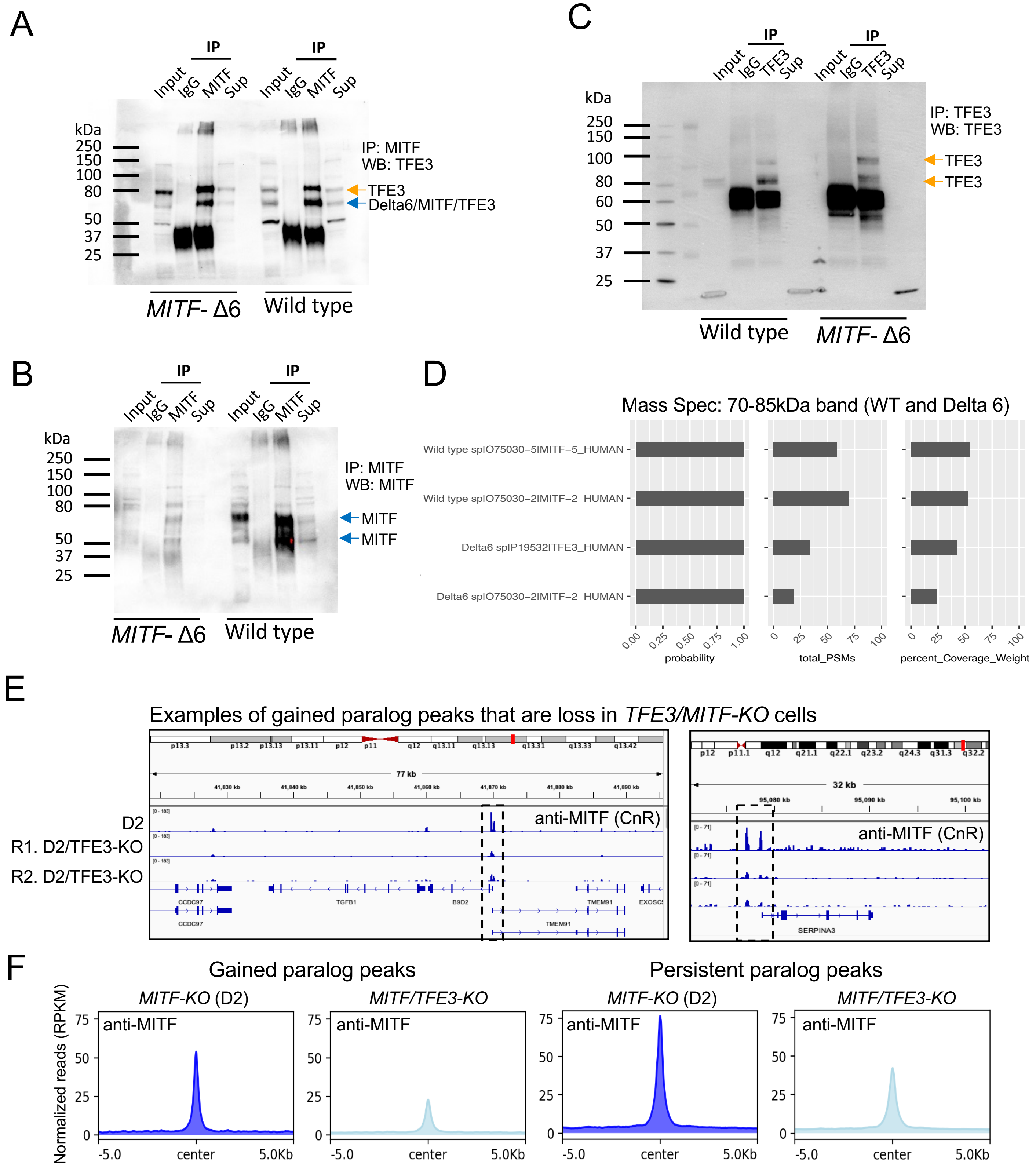
A
IP
Input IgG MITF Sup
kDa
250
150
100
80
50
37
25
IP: MITF
WB: TFE3
TFE3
Delta6/MITF/TFE3
MITF- Δ6
Wild type
B
IP: MITF
WB: MITF
MITF
MITF
C
IP
Input IgG TFE3 Sup
IP: TFE3
WB: TFE3
TFE3
TFE3
60
D
Mass Spec: 70-85kDa band (WT and Delta 6)
Wild type sp|O75030-5|MITF-5_HUMAN
Wild type sp|O75030-2|MITF-2_HUMAN
Delta6 sp|P19532|TFE3_HUMAN
Delta6 sp|O75030-2|MITF-2_HUMAN
probability
total_PSMs
percent_Coverage_Weight
E
Examples of gained paralog peaks that are loss in TFE3/MITF-KO cells
D2
R1. D2/TFE3-KO
R2. D2/TFE3-KO
anti-MITF (CnR)
77 kb
32 kb
CCDC97
TGFB1
B9D2
TMEM91
SERPINA3
F
Gained paralog peaks
Persistent paralog peaks
MITF-KO (D2)
MITF/TFE3-KO
anti-MITF
Normalized reads (RPKM)
-5.0
center
5.0Kb

Figure S4

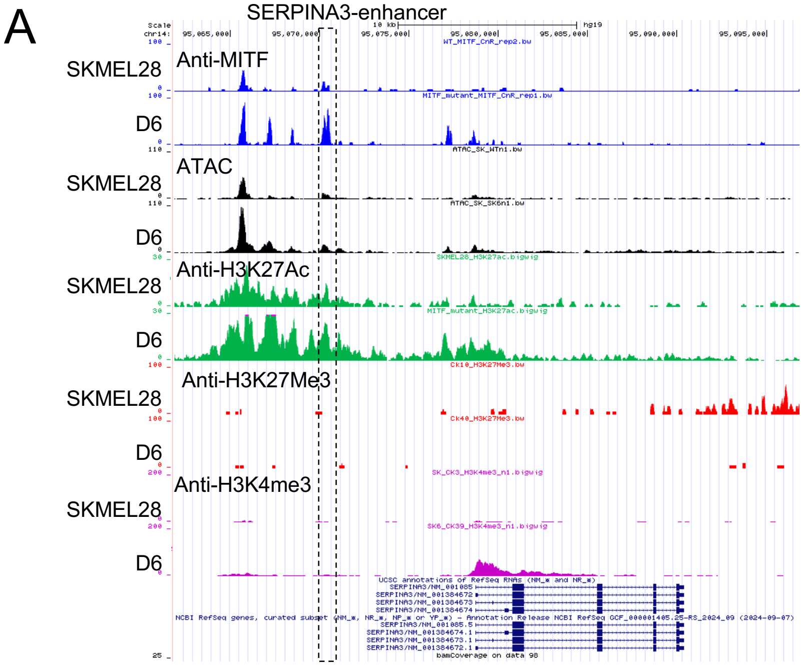


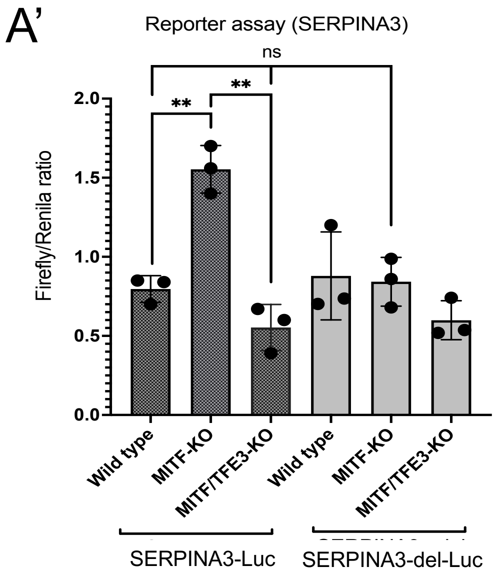


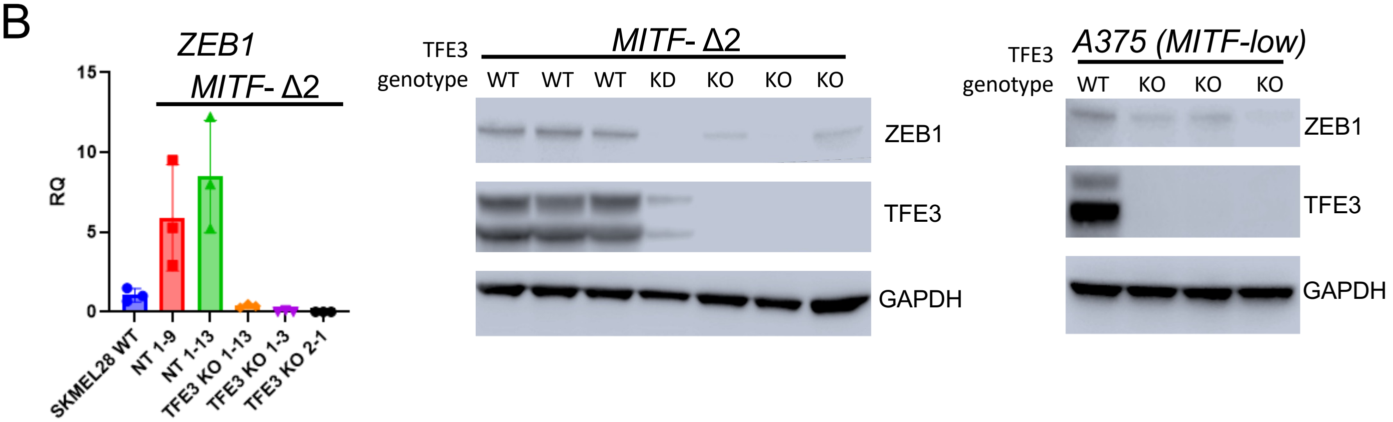


C

Gene Ontology (GO) analysis: TFE3-dependent

| GO | # of genes | FDR |
|---|---|---|
| Cell differentiation | 132 | $2.58 \times 10^{-07}$ |
| Angiogenesis | 49 | $2.9 \times 10^{-05}$ |
| Cell motility | 46 | $3.78 \times 10^{-03}$ |
| Cell migration | 33 | $4.0 \times 10^{-03}$ |

D

Gene Ontology (GO) analysis: TFE3-independent

| GO | # of genes | FDR |
|---|---|---|
| Multicellular organismal process | 506 | $2.7.1 \times 10^{-21}$ |
| Extracellular matrix organization | 77 | $6.5 \times 10^{-18}$ |
| Regulation of type B pancreatic cell proliferation | 33 | $2.8 \times 10^{-04}$ |
| Lymphocyte mediated immunity | 20 | $3.8 \times 10^{-02}$ |

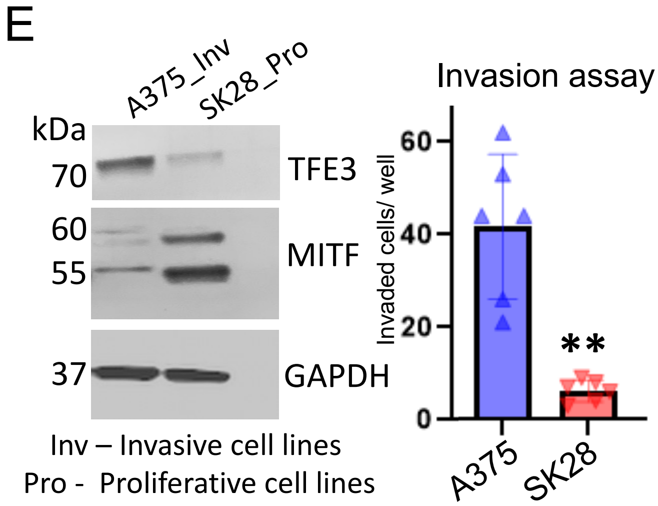


Inv – Invasive cell lines
Pro - Proliferative cell lines

**Figure S5**

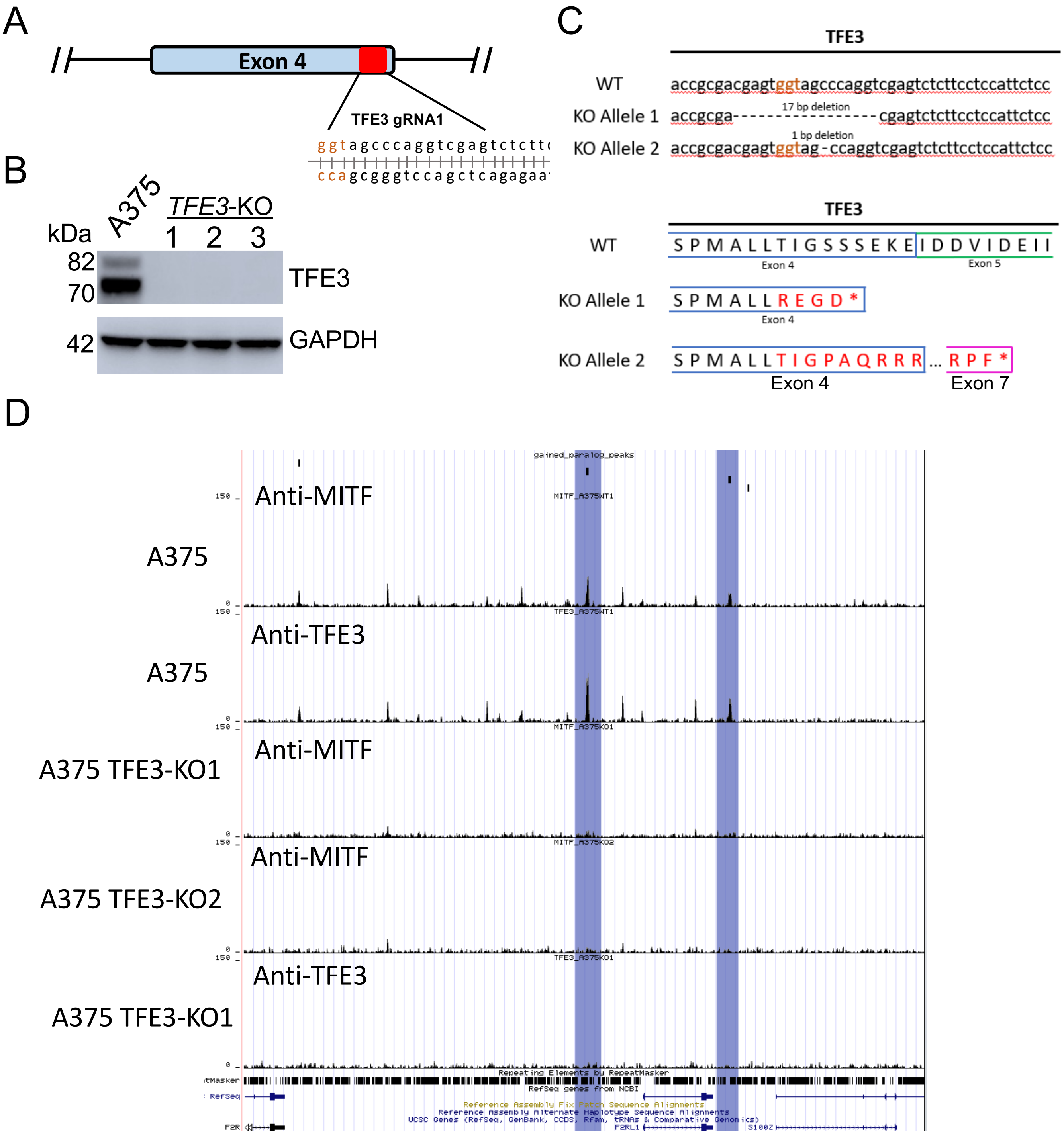
A
Exon 4
TFE3 gRNA1
ggtagcccaggtcgagtctctt
ccagcgggtccagctcagagaa
B
A375
TFE3-KO
1 2 3
kDa
82
70
TFE3
42
GAPDH
C
TFE3
WT
accgcgacgagtggtagcccaggtcgagtctcttcctccattctcc
KO Allele 1
accgcga-----------------cgagtctcttcctccattctcc
17 bp deletion
KO Allele 2
accgcgacgagtggtag-ccaggtcgagtctcttcctccattctcc
1 bp deletion
TFE3
WT
S P M A L L T I G S S S E K E I D D V I D E I I
Exon 4
Exon 5
KO Allele 1
S P M A L L R E G D *
Exon 4
KO Allele 2
S P M A L L T I G P A Q R R R ... R P F *
Exon 4
Exon 7
D
Anti-MITF
A375
Anti-TFE3
A375
Anti-MITF
A375 TFE3-KO1
Anti-MITF
A375 TFE3-KO2
Anti-TFE3
A375 TFE3-KO1
150
0
gained_paralog_peaks
MITF_A375WT1
TFE3_A375WT1
MITF_A375KO1
MITF_A375KO2
TFE3_A375KO1
Repeating Elements by RepeatMasker
RefSeq genes from NCBI
Reference Assembly Fix Patch Sequence Alignments
Reference Assembly Alternate Haplotype Sequence Alignments
UCSC Genes (RefSeq, GenBank, CCDS, Rfam, tRNAs & Comparative Genomics)
RefSeq
F2R
F2RL1
S100Z

**Figure S6**

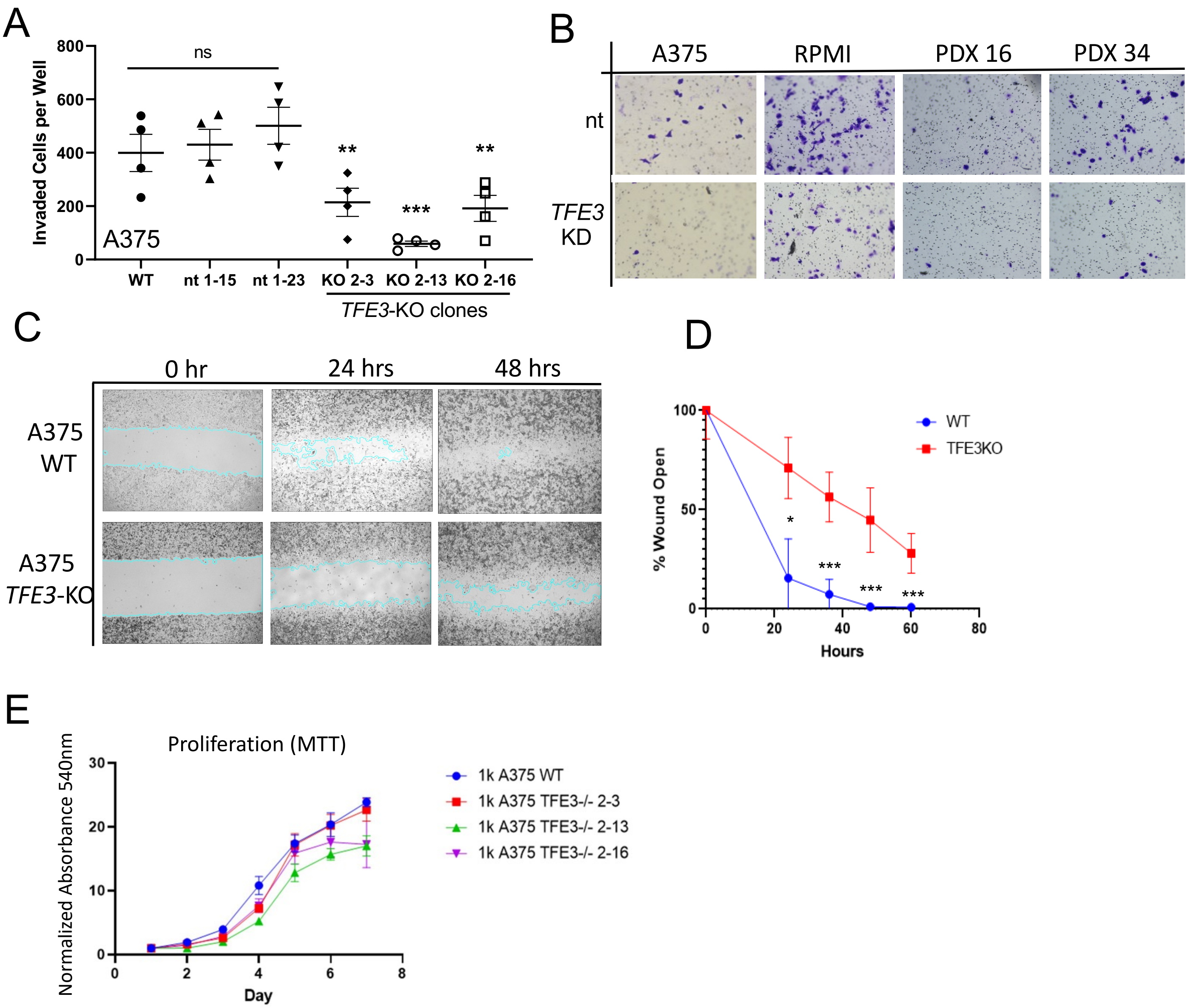

A
Invaded Cells per Well
ns
A375
WT
nt 1-15
nt 1-23
KO 2-3
KO 2-13
KO 2-16
TFE3-KO clones
B
A375
RPMI
PDX 16
PDX 34
nt
TFE3
KD
C
0 hr
24 hrs
48 hrs
A375
WT
A375
TFE3-KO
D
% Wound Open
Hours
WT
TFE3KO
E
Proliferation (MTT)
Normalized Absorbance 540nm
Day
1k A375 WT
1k A375 TFE3-/- 2-3
1k A375 TFE3-/- 2-13
1k A375 TFE3-/- 2-16


**Figure S7**

A

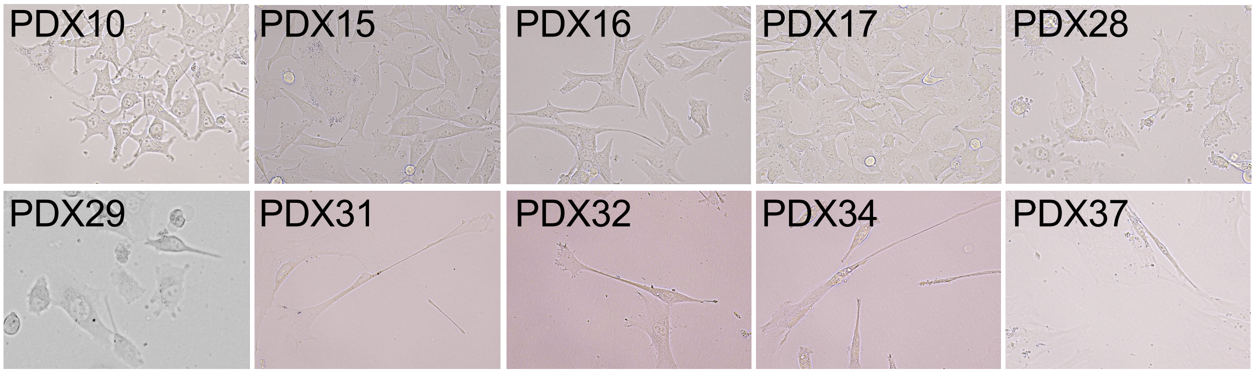
PDX10
PDX15
PDX16
PDX17
PDX28
PDX29
PDX31
PDX32
PDX34
PDX37

**Figure S8**

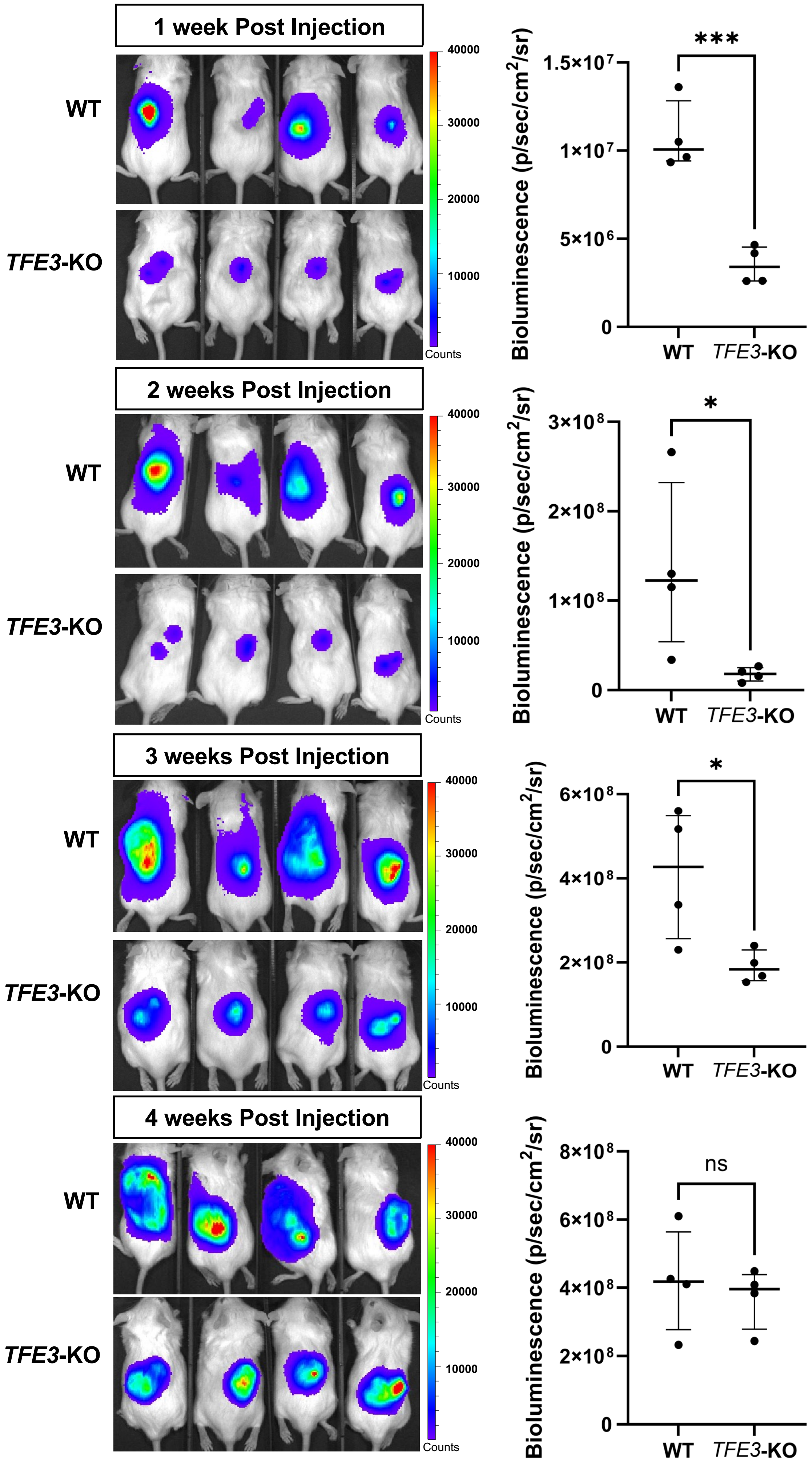

1 week Post Injection
WT
TFE3-KO
40000
30000
20000
10000
Counts
Bioluminescence (p/sec/cm2/sr)
1.5×10^7
1×10^7
5×10^6
0
***
WT
TFE3-KO
2 weeks Post Injection
WT
TFE3-KO
40000
30000
20000
10000
Counts
Bioluminescence (p/sec/cm2/sr)
3×10^8
2×10^8
1×10^8
0
*
WT
TFE3-KO
3 weeks Post Injection
WT
TFE3-KO
40000
30000
20000
10000
Counts
Bioluminescence (p/sec/cm2/sr)
6×10^8
4×10^8
2×10^8
0
*
WT
TFE3-KO
4 weeks Post Injection
WT
TFE3-KO
40000
30000
20000
10000
Counts
Bioluminescence (p/sec/cm2/sr)
8×10^8
6×10^8
4×10^8
2×10^8
0
ns
WT
TFE3-KO


**Figure S9**

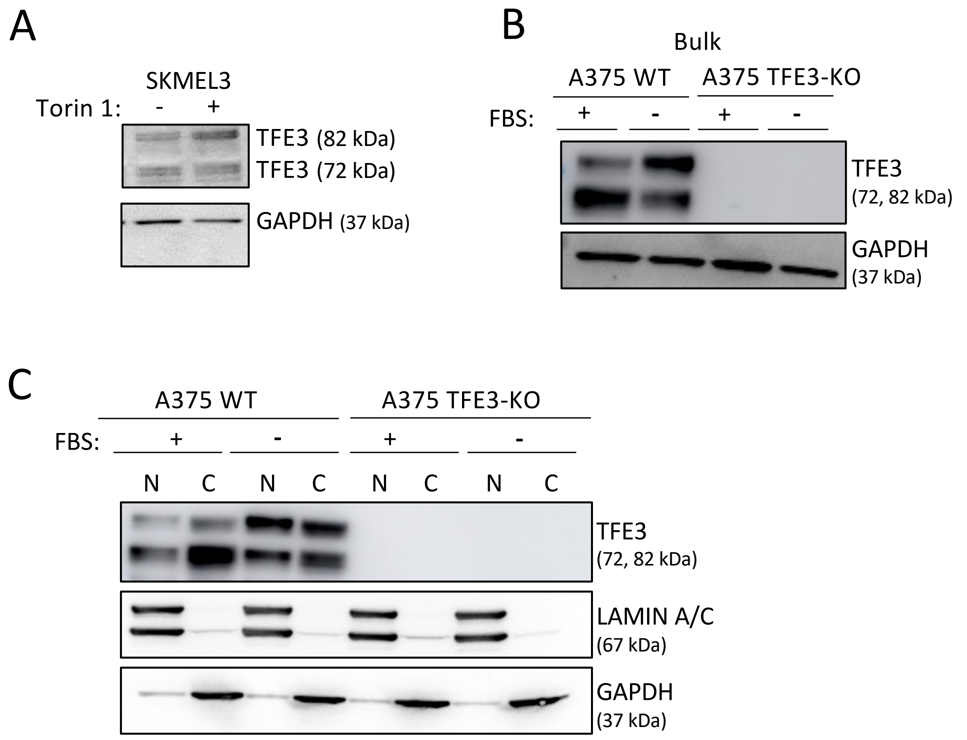
A
SKMEL3
Torin 1: - +
TFE3 (82 kDa)
TFE3 (72 kDa)
GAPDH (37 kDa)
B
Bulk
A375 WT
A375 TFE3-KO
FBS: + - + -
TFE3
(72, 82 kDa)
GAPDH
(37 kDa)
C
A375 WT
A375 TFE3-KO
FBS: + - + -
N C N C N C N C
TFE3
(72, 82 kDa)
LAMIN A/C
(67 kDa)
GAPDH
(37 kDa)

**Figure S10**

A

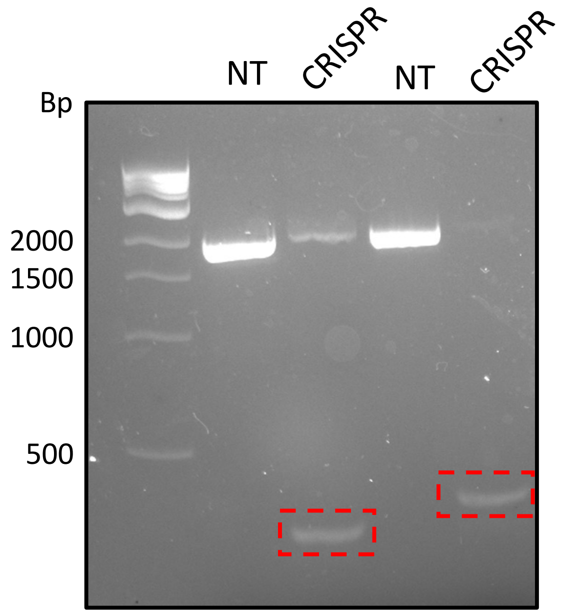


NT = Non-targeting

CRISPR = Double gRNA approach to KO FNIP2

Sequence confirmed cut DNA fragments (*FNIP2*)

B

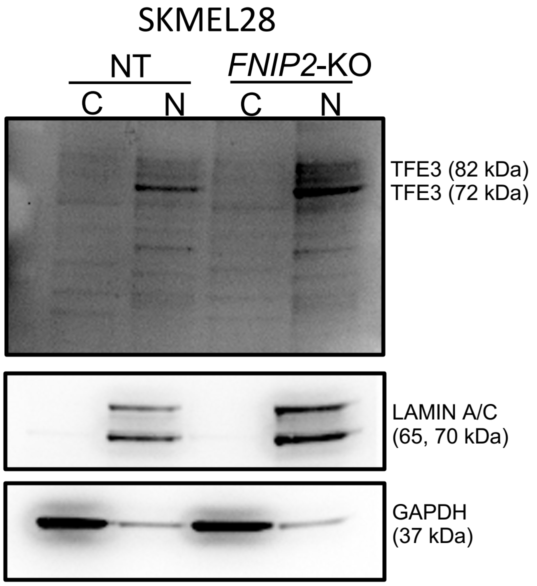


**Figure S11**

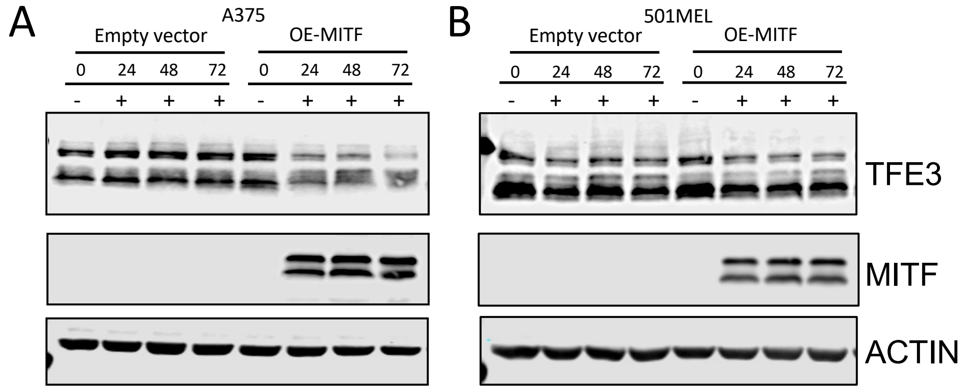
A
A375
Empty vector
OE-MITF
0 24 48 72 0 24 48 72
- + + + - + + +
B
501MEL
Empty vector
OE-MITF
0 24 48 72 0 24 48 72
- + + + - + + +
TFE3
MITF
ACTIN

**Figure S12**

**Supplemental Tables:**

**Table S1:** Differential gene expression and anti-MITF and anti-TFE3 target genes in SKMEL28 and *MITF*-KO (D2, D6) cell lines.

**Table S2:** Differential gene expression comparing double *MITF/TFE3*-KO (clone 13) and *MITF*-KO (D2) cell lines.

**Table S3:** Differential gene expression comparing double *MITF/TFE3*-KO (clone 3) and *MITF*-KO (D2) cell lines.

**Table S4:** Upregulated genes in *MITF*-KO cells vs. SKMEL28 cells that are associated with naïve human embryonic stem cell identity.

**Table S5:** Differential gene expression comparing *TFE3*-KO and A375 cell lines.

**Table S6:** Downregulated genes in *MITF*-KO cells vs. SKMEL28 cells within the mTORC1 signaling pathway.

**Supplemental Figures:**

**Figure S1: Multiple sequence alignment of members of the MIT/TFE transcription factor family and the anti-MITF antibody immunogen sequence**. Amino acid sequences of MITF, TFE3, TFEB, and TFEC, recovered from UniProt and aligned to the immunogen sequence of the anti-MITF (HPA003259, Sigma) using CLUSTALW [53].

**Figure S2: Overlap of anti-MITF CUT&RUN peaks in D2 and D6 cell lines.**

(**A**) Venn diagram illustrating the overlap of anti-MITF CUT&RUN-seq peaks identified by MACS2 in the *MITF-KO* D2 and D6 cell lines.
(**B-C**) Pearson correlation coefficient of true-MITF peaks and gained-paralog peaks of replicate anti-MITF CUT&RUN experiments in SKMEL28 and *MITF*-KO (D6) cell lines.

**Figure S3: Characterization of anti-MITF CUT&RUN peaks in melanoma cells.**

(**A**) Density heatmap representing anti-MITF and anti-H3K27Ac CUT&RUN-Seq, as well as ATAC-Seq, in *MITF-WT* and *MITF-KO* (D6) SKMEL28 cell lines. Peak regions represent true MITF peaks, gained-paralog peaks, and persistent paralog peaks, as indicated.

(**B**) Pie charts showing percentage of overlap of true MITF peaks, gained-paralog peaks, and persistent paralog peaks with MITF-regulated enhancer subtypes, as labeled.

(**C, D**) Gene Ontology (GO) analysis of genes occupied by (**D**) true MITF peaks and (**E**) gained-TFE3 peaks (peaks are within 100 kb of a gene TSS).

**Figure S4: Cross-reactivity of anti-MITF antibodies with TFE3.**

(**A**) Immunoblotting for TFE3 in anti-MITF immunoprecipitates from lysates of *MITF-WT* and *MITF-KO* (D6) cells. A band corresponding to the expected molecular weight of TFE3 (~82 kDa) is labeled yellow, and a band corresponding to the delta6 variant, wild type MITF or short length isoform of TFE3 labeled blue.

(**B**) Immunoblotting for MITF in the samples analyzed in A. Bands at the expected molecular weight of MITF (~60 kDa) are labeled.

(**C**) Immunoblotting of anti-TFE3 immunoprecipitated from lysates of *MITF-WT* and *MITF-KO* SKMEL28 cells. Two distinct bands at the expected molecular weight for TFE3 (~72, 82 kDa) are labeled.

(**D**) Mass spectrometry analysis of anti-MITF immunoprecipitates from lysates of *MITF-WT* immunoprecipitation using an anti-MITF antibody, samples as labeled.

(**E**) Screenshots of IGV (hg19) tracks representing anti-MITF CUT&RUN in *MITF*-KO (D2) and two clones of *MITF/TFE3*-KO (D2/*TFE3*-KO) cell lines at gained paralog peaks near TGFB and SERPINA3 (dashed box).

(**F**) Plot profile representing anti-MITF CUT&RUN signal in *MITF*-KO (D2) and *MITF/TFE3*-KO (D2/*TFE3*-KO) SKMEL28 cell lines. Normalized reads in RPKM. Peak center and +/- 5kb as shown. Peak regions represent loci occupied by gained paralog peaks (left) in D2 cells and persistent paralog peaks (right) in SKMEL28 and *MITF-KO* D2 and D6) cells.

**Figure S5: TFE3 is a transcriptional activator in *MITF-KO* (D2 and D6) cell lines.**

(**A**) Histogram demonstrating activity of the SERPINA3-e1 and SERPINA3-del reporter in SKMEL28, *MITF*-KO and double *MITF/TFE3*-KO cell lines. Individual dots represent biological experiments (n=3). Students t-test; **P-value <0.01, ***P-value <0.001.

(**B**) (Left) qPCR-based quantitation of relative ZEB1 expression in *MITF-WT*, *MITF-KO* (D2), and *MITF-KO* (D2)/*TFE3-KO* SKMEL28 cells. (Middle and right) Immunoblotting for ZEB1 and TFE3 in *MITF-KO* (D2) vs. *MITF-KO* (D2)/*TFE3-KO* cells, and *TFE3-WT* vs *TFE3-KO* cells. *GAPDH* served as a loading control.

(**C**) Gene Ontology (GO) terms identified for genes with higher expression in *MITF-KO* cells than in *MITF-WT* cells.

(**D**) Gene Ontology (GO) terms identified for genes with higher expression in *MITF-WT* cells than in *MITF-KO* cells.

(**E**) Immunoblotting for TFE3 and MITF in *TFE3-WT* and *TFE3-WT* A375 (invasive) and *MITF-WT* SKMEL28 (proliferative) cells. GAPDH is utilized as loading control. (**E**) Quantitation of invasiveness of *TFE3-WT* A375 vs. *MITF-WT* SKMEL28 cells, with number of cells that crossed a Matrigel coated Boydan chamber plotted of the y-axis. Students t-test; **P-value <0.01.

**Figure S6: Isolation of *TFE3-KO* clonal cell lines and anti-MITF CUT&RUN-seq.**

(**A**) Schematic of CRISPR/Cas9 gRNA target region used for TFE3 knockout.

(**B**) Immunoblotting for TFE3 in *TFE3-WT* A375 cells vs. individually isolated *TFE3-KO* A375 clones. GAPDH served as a loading control.

(**C**) Alignments of (top) sequences of *TFE3* gene in *TFE3-WT* and *TFE3-WT* KO A375 cell lines, and (bottom) of associated translated amino acid sequences. Red text indicates frameshift of AA sequence.

(**D**) Screenshot of UCSC (hg19) genome browser visualization of bigwig files generated from anti-MITF and anti-TFE3 CUT&RUN in A375 and TFE3-KO cell lines at the F2RL1 locus. Blue highlight represents a gained-TFE3 in D6 and D2 *MITF*-KO cell lines.

**Figure S7: Cell biological endpoint assays in *TFE3-KO* cells**

(**A**) Quantitation of results for Transwell invasion assay for *TFE3-WT* A375 cells and CRISPR clones that did not target TFE3 vs. CRISPR *TFE3-KO* clones. Individual dots represent biological experiments (n=4) with three technical replicates. **P-value <0.01, ***P-value <0.001.

(**B**) Representative 10X brightfield images taken during Transwell invasion assay, in untreated (nt) and KD cells: A375 (*TFE3-WT*), RPMI-7951, PDX 16, and PDX 34.

(**C-D**) Representative 10X brightfield images taken during scratch wound-healing assay of *TFE3-WT* A375 and *TFE3-KO* cells. (Left) 4X brightfield images taken at 0, 24, and 48 hours post scratching. (**D**) Quantification of open wounds, using ImageJ. Individual dots represent biological experiments (n=3) with three technical replicates. *P-value<0.05, ***P-value<0.001.

(**E**) Quantification of cell proliferation assay for *TFE3-WT* cells vs *TFE3-KO* A375 clones, based on normalized absorbance at 540nm. Individual dots represent biological experiments (n=6) with three technical replicates.

**Figure S7: Brightfield images of PDX cell lines.**

**Figure S9: Imaging of tumor intradermal injections at 1-4 weeks.**

*In vivo* imaging of primary tumors derived from intradermal injection of luciferase-expressing A375 (n=4) and *TFE3*-KO (n =4) into immunocompromised NOD/SCID mice 1 – 4 weeks post-injection. Imaging was performed with a 60-second exposure, 10 minutes after intraperitoneally injection of firefly luciferin (120 mg/kg). Student's t-test; * P-value <0.05.

(**A'-B'**) Quantification of bioluminescence from xenografts generated by TFE3-WT and TFE3-KO A375 cells at (A') 1 week and (B') 3 weeks post-injection (N=4).

**Figure S10: TFE3 response to serum depletion in A375 cell lines.**

(**A**) Immunoblotting of anti-TFE3 in lysates of SKMEL3 cell lines, +/- Torin1 (1 uM) or control DMSO treatment for 24 hours.

(**B**) Immunoblotting of anti-TFE3 in lysates of A375 and *TFE3*-KO cell lines, +/- fetal bovine serum (FBS) for 24 hours.

(**C**) Immunoblotting of anti-TFE3 in cytoplasmic and nuclear lysates of A375 and *TFE3*-KO cell lines, +/- fetal bovine serum (FBS) for 24 hours.

**Figure S11: Knockout of FNIP2 from MITF-high SKMEL28 cell lines.**

(**A**) Agarose gel image of DNA fragments amplified by PCR using primers flanking exons 4 and 6 of FNIP2 (NM_001323916). gRNAs were designed to target an 1800 bp segment within FNIP2 containing exons 4, 5, and 6. Primers flanking these gRNA target sites produce a 2000 bp DNA band in the presence of the full-length region, while a successfully edited DNA product would produce a shorter ~200 bp band, indicated by the red dashed box. A full-length band in CRISPR samples may indicate single guide editing or duel editing that does not remove the middle segment.

(**B**) Immunoblotting for TFE3 of cytoplasmic (C) and nuclear (N) fractions of lysates from WT and FNIP2 edited SKMEL28 cells.

**Figure S12: Overexpression of MITF in A375 and 501MEL cell lines.**

(**A-B**) Immunostaining for anti-TFE3 and anti-HA-MITF in stable doxycycline inducible (A) A375 and (B) 501MEL cell lines along with empty vector controls. Cells were treated with 0.1 mg/mL doxycycline for 0, 24, 48, and 72 hours.
